# Carbon Dioxide Controls Fungal Fitness and Skin Tropism of *Candida auris*

**DOI:** 10.1101/2024.04.12.589292

**Authors:** Trinh Phan-Canh, Philipp Penninger, Saskia Seiser, Narakorn Khunweeraphong, Doris Moser, Tamires Bitencourt, Hossein Arzani, Weiqiang Chen, Lisa-Maria Zenz, Andrej Knarr, Diana Cerbu, Sabrina Jenull, Christoph Müller, Michaela Lackner, Giuseppe Ianiri, Anuradha Chowdhary, Markus Hartl, Adelheid Elbe-Bürger, Karl Kuchler

## Abstract

The pronounced skin tropism and pan-antifungal resistance traits of the fungal pathogen *Candida auris* stand out as a serious health threat. Here, we show that a carbonic sensing pathway (CSP) promotes development of resistance to amphotericin B through a reactive oxygen species (ROS) response, as well as ectopic cell wall and membrane lipid homeostasis. Mechanistically, the transcription factor Rca1 acts in cooperation with Efg1 to control the expression and activity of the carbonic anhydrase Nce103 as a key effector component. The conversion of carbon dioxide to bicarbonate provides a direct link to energy metabolism, facilitating colonization and growth on skin tissues. Native mouse and human skin models unequivocally show that the CSP is essential for maintaining skin tropism as well as fungal fitness. Curiously, upon ablation of Rca1 and Efg1, *C. auris* debilitates efficient growth on native skin. Collectively, our findings highlight critical roles of the CSP in *C. auris* skin tropism and antifungal drug resistance. The work suggests therapeutic options for disrupting skin colonization and thus preventing infections.

**Highlights:**

✓ Proteo-transcriptomics links a carbonic sensing pathway (CSP) to *C. auris* multidrug resistance
✓ The Nce103 carbonic anhydrase controls drug resistance as a key component of the CSP
✓ The transcription factors Rca1 and Efg1 control Nce103 and link CSP with *C. auris* skin tropism
✓ CSP acts through ectopic ROS response, cell wall architecture and membrane lipid function
✓ CSP is required for *C. auris* fitness and efficient growth and colonization of skin tissues

**Result contents:**

✓ Integrated omics reveals multidrug-resistant mechanisms in *C. auris*
✓ CO_2_-sensing controls amphotericin B resistance (AMB^R^) traits through Rca1 and Efg1
✓ The carbonic anhydrase Nce103 governs susceptibility to amphotericin B
✓ The CSP influences AMB^R^ by maintaining reactive oxygen species homeostasis
✓ The CSP controls AMB^R^ via cell membrane and cell wall remodelling
✓ The CSP regulates fungal fitness through controlling energy metabolism
✓ *C. auris* requires the CSP for skin colonization

## 1. Introduction

*Candida auris* is an emerging human fungal pathogen causing disseminated infections of high mortality (30-72%) in individuals with chronic diseases or impaired immunity^1–3^. Importantly, the high rate of multidrug resistance traits^1,2,4,5^ pose a serious challenge to clinical therapy with the available antifungal arsenal. This is further exacerbated by the pronounced skin colonization and the resulting ease of transmission by skin-to-skin contact in hospital settings, which represents an unsolved medical problem of utmost relevance^6–8^. Indeed, since 2009^9^, *C. auris* has spread to more than 50 countries, causing outbreaks in intensive care units and nursing homes^10–12^. Most importantly, *C. auris* is the first fungal pathogen showing untreatable clinical pan-antifungal multidrug resistance traits to all clinically used antifungal drugs, including azoles (up to 90%), amphotericin B (AMB, up to 50%) and echinocandins (up to 8%), as well as flucytosine^4,5,13,14^. However, the molecular mechanisms of this phenomenon, particularly the high rate of amphotericin B resistance (AMB^R^) remain not fully understood^15–18^. Hence, we employed advanced integrated proteo-transcriptomics datasets and genetic tools to uncover novel functions of a carbonic sensing pathway in regulating AMB^R^, skin tropism and virulence of *C. auris*.

## 2. Results

### 2.1. Integrated omics reveals multidrug-resistant mechanisms in *C. auris*

Transcriptomics of multidrug-resistant (MDR) versus drug-sensitive *C. auris* clinical isolates revealed numerous genes of carboxylic acid metabolism, mitochondrial function, translation, as well as membrane transporters as critical indicators of antifungal resistance in *C. auris*^19–21^. Hence, we integrated transcriptomics datasets with proteomics data from the same strains to identify major genes/pathways implicated in *C. auris* MDR traits. We subjected whole cell extracts from the pair of resistant (462/P/14 – R1) and sensitive (2431/P/16 – S) *C. auris* isolates^19,21^ (Fig. 1a, b, and Extended Data Fig. 1a, b) to shotgun proteomics. Differentially abundant proteins represent various metabolic processes, including amino acid, nucleic acid and lipid biogenesis (Extended Data Fig. 1c, d, e). Remarkably, most membrane transporter families are highly enriched in resistant *C. auris* strains (Extended Data Fig. 1e, f), including the Mdr1 major facilitator and the Cdr1 multidrug efflux ABC transporter responsible for azole resistance^20,22–28^. In addition, the isolates R1 and 1133/P/13R (R2), but not S, carried the mutational change S639F in the Fks1 glucan synthase (Extended Data Fig. 1g)^19,29^, thus explaining in part echinocandin resistance in R1 and R2^22^. Of note, about 50% of all clinical *C. auris* isolates are resistant to AMB^7,14^, but mechanistic explanations are just emerging^13,15,22^.

**Fig. 1.**
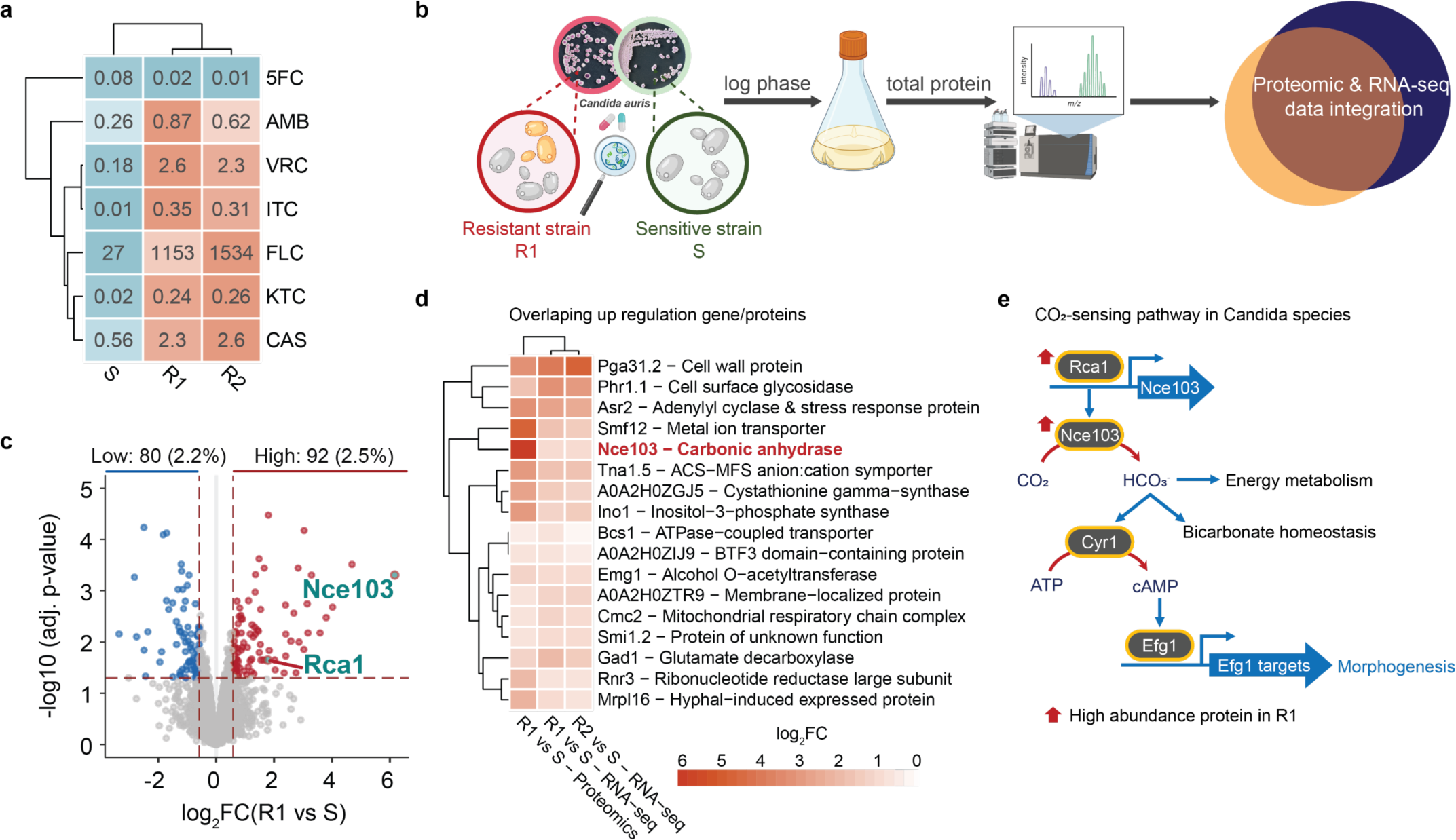
│ A CO_2_-sensing pathway is linked to drug resistance. **a**, Heatmap showing inhibitory concentration IC_50_ values for pan-drug-resistant (R1, R2) and drug-sensitive (S) clinical strains after dose-response assays at 30 °C for 30 hours in RPMI-MOPS-pH7 (RPMI). 5FC: 5-flourocytosine, AMB: amphotericin B, VRC: voriconazole, ITC: itraconazole, FLC: fluconazole, KTC: ketoconazole, CAS: caspofungin. **b**, Cartoon illustrates the proteomic workflow and data analysis, drawing with BioRender. **c**, Volcano plot of proteomics data shows differentially abundant proteins (DAPs) between resistant and sensitive strains based on LIMMA analysis. **d**, Heatmap indicates 17 proteins/genes of high abundance in resistant strains from both proteomics and RNA-seq datasets. **e**, Scheme reveals a CO_2_-sensing pathway (CSP) controlling a link between energy metabolism and morphogenetic switching. Differential expression analysis filters genes/proteins with log_2_FC ± 0.58 and adjusted p value < 0.05 (calculated with Benjamini-Hochberg – BH).

The overlay of proteomics and RNA sequencing (RNA-seq) datasets revealed the top 5 high abundance proteins, including homologues of the cell wall components Pga31.2 (B9J08_004476) and Phr1 (B9J08_000918), the adenylyl cyclase Asr2 (B9J08_000849) and the manganese transporter Smf12 (B9J08_002789). Most strikingly, the carbonic anhydrase Nce103 (B9J08_000363) that converts CO_2_ into bicarbonate HCO ^-^ was the highest abundance protein in R1 (Fig. 1d, Extended Data Fig. 1h). In addition, the transcription factor Rca1, a key regulator of carbonic anhydrase Nce103, was also enriched in strain R1 (Fig. 1c). Of note, despite numerous attempts, we failed to ablate *NCE103*, even when supplementing 5.5% CO_2_, suggesting that *NCE103* is an essential gene in *C. auris*. The integrated data indicate a putative CO_2_-sensing pathway (CSP) encompassing the Rca1-regulated upstream Nce103 carbonic anhydrase (Fig. 1e). Remarkably, bicarbonate (HCO_3_ ^-^) plays critical roles in energy metabolism and morphogenesis, since it is acting through the cAMP/PKA signalling pathway that converges at the regulator Efg1^30^, which is fully consistent with metabolic processes suggested by the enrichment analysis (Extended Data Fig. 1c, d).

### 2.2. CO_2_-sensing controls AMB^R^ traits through Rca1 and Efg1

While CO_2_-sensing has been reported in some microbial pathogens^30–35^, possible functions in drug resistance varies among pathogens^30,32^, perhaps owing to distinct wiring of regulatory networks^36^. Importantly, orthologues of the carbonic anhydrase regulator Rca1 are only found in Candida and *Saccharomyces* spp. (Fig. 2a). Interestingly, Rca1 and the highly conserved HTH APSES-type regulator Efg1 (Extended Data Fig. 2) appear to cooperate for CO_2_ conversion during adaptation of *C. albicans* to different host niches^37,38^.

**Fig. 2.**
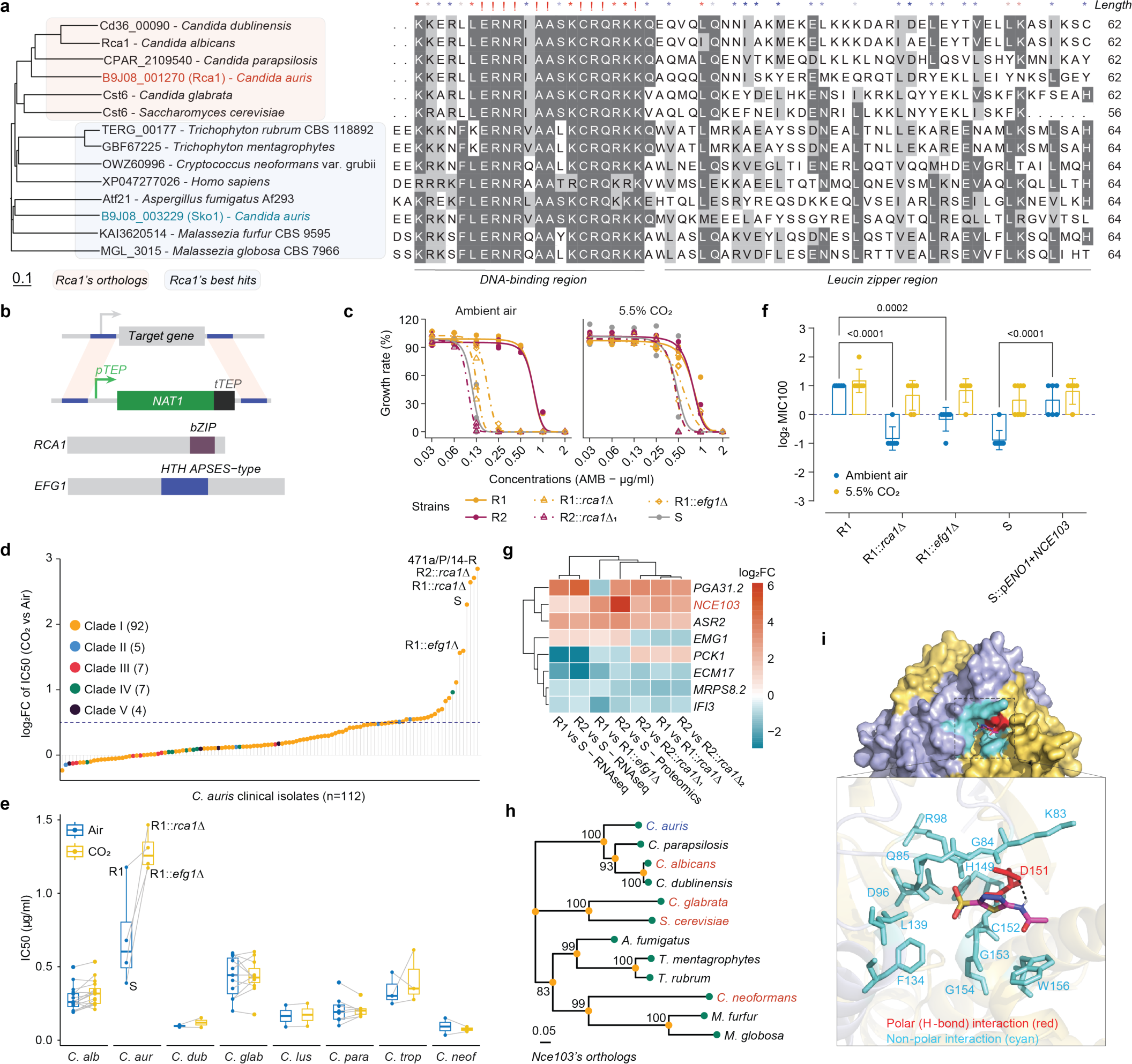
│ A CSP controls antifungal drug resistance. **a**, Multiple sequence alignments reveal the evolutionary conservation of the bZip domain in Rca1. **b**, Gene deletion of *RCA1* and *EFG1*. **c-f**, Dose-response assay shows growth inhibition of amphotericin B (AMB) in ambient air and 5.5% CO_2_. Culture conditions were RPMI at 37 °C for 30 hours. **c**, AMB dose-response curves for strains R1, R2, S, *rca1*Δ and *efg1*Δ mutants. **d**, log_2_FC of IC_50_ values of 115 *C. auris* clinical isolates representing five clades (colored dots) in 5.5% CO_2_ versus ambient air with *rca1*Δ and *efg1*Δ as controls. **e**, IC_50_ values of different clinical isolates of *C. albicans (C. alb), C. auris (C. aur), C. dubliniensis (C. dub), C. glabrata (C. glab), C. lusitaniae (C. lus), C. parapsilosis (C. para), C. tropicalis (C. trop)* and *Cryptococcus neoformans (C. neof)*. **f**, MIC_100_ values for AMB recorded at 48 hours. Data expressed as log_2_. Two-way ANOVA with Tukey’s multiple comparisons test was applied. **g**, Heatmap depicts 8 major Rca1 and Efg1 target genes. **h**, Neighbor-joining phylogenetic tree analysis of fungal Nce103 homologues (bootstrap ∼ 1000 times). Numbers indicate bootstrap values (%). **i**, Homology model of *C. auris* Nce103 reveals conservation of binding pocket for the ACZ inhibitor. Polar (red) and non-polar (cyan) interacting residues based on Coccomyxa beta-carbonic anhydrase in complex with acetazolamide (PDB: 3UCJ). Scale bar in **(a, g)** indicates the number of differences between sequences.

To uncover how the CSP controls drug resistance of *C. auris*, we deleted *RCA1* and *EFG1* in resistant clinical strains (Fig. 2b). Remarkably, ablation of *RCA1* or *EFG1* re-sensitized otherwise resistant *C. auris* isolates to AMB, with 4-8-fold decreases of IC_50_ values (Fig. 2c, Extended Data Fig. 3a, b). Moreover, *rca1*Δ and *efg1*Δ mutants showed slightly increased susceptibility to 5-fluorocytosine (5FC) although not to caspofungin (CAS) or voriconazole (VRC) (Extended Data Fig. 3a, c). To verify the role of the CSP, we repeated dose-response experiments in the presence and absence of 5.5% CO_2_ to mimic the distinct CO_2_ concentrations on skin surface or in deeper tissues. Intriguingly, *rca1*Δ, *efg1*Δ, and even the sensitive strain S fully restored AMB^R^ in the presence of 5.5% CO_2_ (Fig. 2c, f) as opposed to AMB sensitive (AMB^S^) in ambient air (∼0.04% CO_2_). Because of pronounced phenotypic variabilities in *C. auris* isolates, we subjected more than 100 clinical isolates from different clades to AMB dose-response assays. Remarkably, CO_2_ significantly enhanced IC_50_ values by 2-8-fold increases in some clinical isolates, which was similar to *rca1*Δ and *efg1*Δ (Fig. 2d). Using a log_2_FC > 0.5 cut-off, some 22 out of 112 strains from different clades (roughly 20%) significantly enhanced IC_50_ values in CO_2_. MIC_100_ values were also recorded after 48 hours to correct for a possible growth bias of distinct isolates, yielding a similar number of clinical isolates (17/112 strains) with increased AMB^R^ in 5.5% CO_2_ (Extended Data Fig. 3d). Of note, AMB dose-response assays for clinical isolates representing other fungal pathogens, revealed that a CO_2_-dependent response engaging Rca1 and Efg1 exists only in *C. auris* (Fig. 2e).

### 2.3. The carbonic anhydrase Nce103 governs susceptibility to AMB

Further, we aimed to clarify the role of Nce103 in AMB^R^. Noteworthy, Nce103 was upregulated in all omics data sets when we compared resistant (R1, R2) to sensitive *C. auris* isolates (S, *rca1*Δ, *efg1*Δ) (Fig. 2g, Supplementary Table 6)^21^. Because Nce103 is essential, we used acetazolamide (ACZ), a specific inhibitor of carbonic anhydrases^39^, to test for consequences after blocking Nce103. Although, we observed a minor inhibitory effect on the growth of *C. auris*, ACZ inhibited *rca1*Δ, *efg1*Δ and strain S to a higher extent (Extended Data Fig. 3e). Of note, R1 was resistant to ACZ under the conditions used, suggesting that Nce103 expression and protein levels could be critical for the extent of drug resistance traits. Indeed, Nce103 levels in R1 were more than 70-fold higher than in the S isolate (Fig. 2g, Supplementary Table 6). Likewise, pairwise comparisons of *NCE103* mRNAs between AMB^R^ and AMB^S^ strains indicated 3-8 times higher expression in AMB^R^ strains^21^. To further support this notion, we next generated a variant S strain overexpressing Nce103, resulting in a 2-4-fold increase of AMB MIC_100_ values (Fig. 2f).

Interestingly, key motifs Nce103 are highly conserved in various fungal pathogens (Fig. 2h; Extended Data Fig. 4a). To confirm that ACZ blocks *C. auris* Nce103, we generated structural homology models (Extended Data Fig. 4b) using the coordinates of the Coccomyxa β-carbonic anhydrase (CA) co-crystallized with ACZ (PDB: 3UCJ^39^). Indeed, ACZ binding pockets are highly conserved in β-CA from algae to fungi (Extended Data Fig. 4c, d). Docking of ACZ to the putative binding region identified in the Nce103 homology model (Extended Data Fig. 4e) revealed that both ACZ and the derivative ethoxzolamide (ETZ) shared common interacting residues (Fig. 2i, Extended Data Fig. 4f). This data confirmed a conserved core structure of β-CA Nce103 across fungal pathogens (Fig. 2i) and supports the observed inhibition of *C. auris* Nce103 by ACZ.

Remarkably, IC_50_ values and the 8-fold increase in 5.5% CO_2_ were similar between strain S and the *rca1*Δ mutants (Fig. 2c, f). To test whether or not sequence variations contribute to this phenomenon, we inspected available RNA-seq datasets^19,21^ for *RCA1, NCE103* and *EFG1* transcripts using the IGV tool^40^. Intriguingly, we discovered a single nucleotide polymorphism (SNP) in *RCA1* at nucleotide position 330, yielding a truncated Rca1 variant with a stop codon replacing Y110 just before the bZIP region (Extended Data Fig. 5a). Indeed, the Y110* Rca1 variant appears unstable and non-functional, since proteomics failed to detect any Rca1-derived peptides from strain S (Extended Data Fig. 5b). Because of the essentiality of Nce103, we hypothesized that sequence variations may be more prevalent in the *RCA1* and *EFG1* loci but not in *NCE103*. Hence, we *blastn*-searched whole-genome shotgun contigs (wgs) and nucleotide collection (nr/nt) NCBI databases on *C. auris*. Strikingly, about ∼20% and ∼10% of 122 *C. auris* clinical isolates harboured SNP variations in *RCA1* and *EFG1,* respectively. Only two rare SNP variations (R214I-K223Q; N224Tfr*9) were found in *NCE103* (2%) across all clades (Extended Data Fig. 5c). Of note, four strains contained frame shift mutations in *RCA1*, leading to a truncated protein, and all were AMB-sensitive^41–44^ (Extended Data Fig. 5d). Hence, genetic variations in *RCA1* and *EFG1* loci may cause varying *NCE103* transcript levels to mount AMB^R^ traits. In summary, Rca1 and Efg1 orchestrate the control AMB^R^ by regulating the levels of Nce103.

### 2.4. The CSP influences AMB^R^ by maintaining reactive oxygen species homeostasis

To address how Nce103 impacts AMB^R^, we exposed both resistant (R) and sensitive (S) strains to dose-response assays with different metabolic inhibitors (Fig. 3a, Extended Data Fig. 6a). Remarkably, S strains displayed increased sensitivity to the mitochondrial electron transport chain (ETC) complex III inhibitor antimycin A, and to the N-linked glycosylation inhibitor tunicamycin (Fig. 3a). This correlated with the dysregulated gene expression after Rca1 or Efg1 disruption (Fig. 3c, Extended Data Fig. 6b). Impairing mitochondrial functions or the accumulation of unfolded proteins in the endoplasmic reticulum can promote accumulation of reactive oxygen species (ROS)^45,46^, thus increasing hydrogen peroxide (H_2_O_2_) susceptibility in *rca1*Δ and *efg1*Δ mutants (Extended Data Fig. 6a). Furthermore, sensitive strains exhibit elevated background levels of intracellular ROS when compared to R1, indicating increased ROS levels in the absence of Rca1 or Efg1 (Fig. 3b, Extended Data Fig. 6c). Indeed, AMB caused significantly higher intracellular ROS in strain S, *rca1*Δ and *efg1*Δ mutants (Fig. 3b, Extended Data Fig. 6c). Increased ROS exacerbates DNA and membrane lipid damage, which was evident in the hypersensitivity to sodium dodecyl sulphate and genotoxic compounds such as methyl methane sulfonate and hydroxy urea (Fig. 3a, Extended Data Fig. 6a). Hence, a dysfunctional CSP impairs mitochondrial as well as ER functions, thereby affecting stress tolerance of *C. auris* (Fig. 3i).

**Fig. 3.**
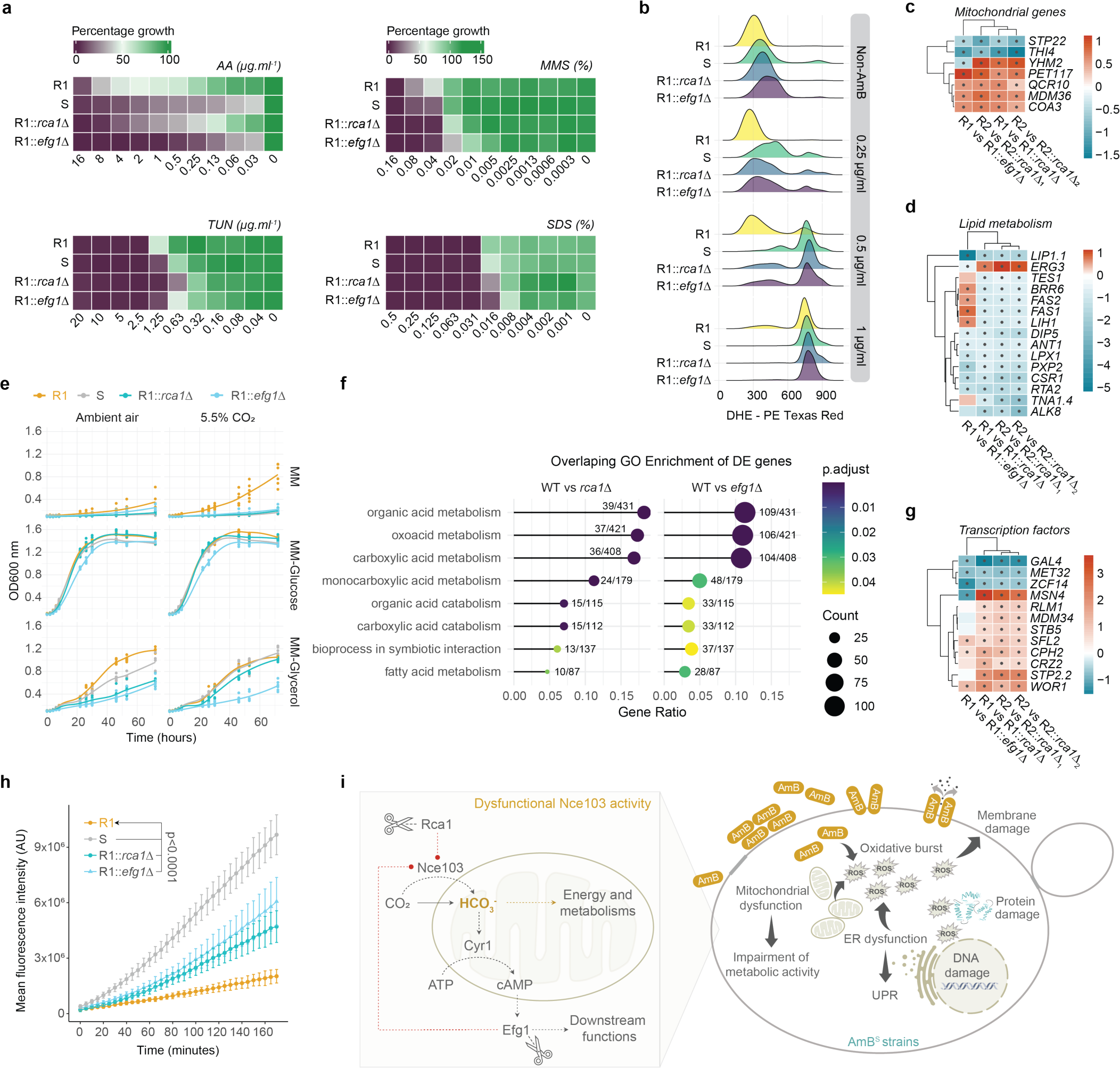
│ Mechanisms implicated in CSP-mediated drug resistance. **a**, Dose-response assays in RPMI for 30 hours at 37 °C with a mitochondrial inhibitor (antimycin A – AA), a protein N-link glycosylation inhibitor (tunicamycin), genotoxic stress agents (methyl methane sulfonate – MMS), a membrane stressor (sodium dodecyl sulphate – SDS). **b**, Fungal cells were exposed to different AMB concentrations for 2 hours, followed by staining with fluorescent dihydroethidium (DHE). Intracellular ROS measured with PE-TexasRed by flow cytometry. **c, d, g**, Heatmaps show differentially expressed genes (DEGs) involved in different cellular processes. Data exported from comparative transcriptomic data of wild-type strains (WT) and independent *rca1*Δ mutants clones. Data for *efg1*Δ were included in the figure after filtered with *rca1*Δ cut-off log_2_FC of ± 0.58. Color scale shows log_2_FC values; black dots indicate adjusted p-value ≤ 0.05. **e**, Growth curves were recorded by OD_600nm_ for 72 hours in minimal medium containing different carbon sources. **f**, Gene ontology (GO) enrichment analysis for DEGs in *rca1*Δ, *efg1*Δ compared to WT. Data were filtered with cut-off log_2_FC of ± 0.58. Shared GO terms between *rca1*Δ and *efg1*Δ were visualized. **h**, Membrane lipid permeability was assessed by a fluorescein diacetate uptake assay. Two-way ANOVA of mean fluorescence intensity (MFI) for time x strains followed by Tukey’s multiple comparison tests using R1 as control group. **i**, Impact of a CSP on AMB stress response and biological pathways.

### 2.5. The CSP controls AMB^R^ via cell membrane and cell wall remodelling

The increased SDS susceptibility hints alterations in membrane lipid functions (Fig. 3a). Hence, we assessed membrane permeability using the lipophilic fluorescent precursor fluorescein diacetate (FDA) as an indicator of protein-independent membrane permeability. Intriguingly, strain S, and the *rca1*Δ and *efg1*Δ mutants showed a substantial increase in FDA membrane permeability (Fig. 3h). This correlated with the dysregulation of 15 lipid metabolism genes in the RNA-seq data of *rca1*Δ and/or *efg1*Δ (Fig. 3d), thus supporting the notion of altered plasma membrane function when the CSP is rendered dysfunctional. Further, expression of *HSP12* gene encoding the unstructured heat shock protein Hsp12 implicated in membrane lipid stress^47^ was significantly impaired (∼1.6-fold decrease in *rca1*Δ, more than 40-fold decrease in *efg1*Δ)(Extended Data Fig. 7a, e), and correlated permeability changes in *efg1Δ* when compared to *rca1Δ* (Fig. 3h). These results suggest that CSP controls membrane lipid function and cell wall architecture through a downstream pathway that engages Efg1 and Hsp12, thereby also affecting antifungal susceptibility. Indeed, comparing wild-type (WT), *rca1*Δ and *efg1*Δ mutants (Extended Data Fig. 8b, e) revealed pronounced changes in the sensitivities to cell wall stressors, including Calcofluor White, Congo Red and Nikkomycin Z (Extended Data Fig. 8a).

In summary, ablation of *RCA1* or *EFG1* in R strains diminished *NCE103* transcript levels and promoted sensitivity to AMB, whereas *NCE103* overexpression in the S strain increased AMB^R^. The CSP controls AMB sensitivity, thus offering a plausible explanation for the exceptionally high polyene resistance rates seen in *C. auris*. Furthermore, the CSP exerts broader effects on key biological functions, including mitochondrial homeostasis, ER-related stress response as well as cell wall and membrane lipid functions, and all appear linked to detrimental AMB-triggered ectopic ROS responses (Fig. 3i).

### 2.6. The CSP regulates fungal fitness through controlling energy metabolism

RNA-seq show that both Rca1 and Efg1 regulate Nce103 levels, which feeds the resulting bicarbonate (HCO ^-^) into metabolic processes^48,49^. Therefore, we overlayed the GO enrichment data of *rca1*Δ and *efg1*Δ mutants, yielding 8 shared biological processes related to carboxylation involving in TCA cycle, as well as amino acid, nucleic acid, carbohydrate and fatty acid metabolism (Fig. 3d, f; Extended Data Fig. 6d). Hence, we inspected growth properties at large in various media. Although amino acid biosynthesis-related transcripts were altered (Extended Data Fig. 6d), elimination of one or all amino acids from synthetic complete (SC) medium had no significant growth impact (Extended Data Fig. 9a). In addition, blocking TOR1 signalling with rapamycin showed no significant differences between S and R strains (Extended Data Fig. 6a). Glucose, not glycerol, partially improved the growth of S strains (*rca1*Δ, *efg1*Δ, S) (Fig. 3e, Extended Data Fig. 9b). Furthermore, *rca1*Δ and *efg1*Δ strains dysregulated the carbon source dependent Gal4 regulator (Fig. 3g), as well as the putative hexose transporter genes *HGT2.1, HGT13.3, HGT19.1, ITR1* (Extended Data Fig. 7d). Although glucose or 5.5% CO_2_ rescued the overall fitness defect, *rca1*Δ strains showed a prolonged lag phase when compared to R1 (Extended Data Fig. 9b, c). In minimal medium with 5.5% CO_2_, strain R1 reached an OD_600nm_ of 1 after 72 hours, whereas *rca1*Δ, *efg1*Δ and strain S failed to grow (Fig. 3e). Moreover, overexpression of *NCE103* in strain S yielded a growth curve similar to R1 (Extended Data Fig. 9d). While the *NCE103* mRNA levels in the overexpression strain were remarkably higher when compared to R1, their growth rates remain comparable (Extended Data Fig. 9d). In conclusion, our data suggest that the CSP establishes an energy-generating mechanism, since it enables *C. auris* to fix even minute levels of CO_2_ when growing in ambient air, as for example on the skin surface.

### 2.7. *C. auris* requires the CSP for skin colonization

In addition to metabolic alterations in S strains, we observed remarkable expression changes of genes linked to cell surface function (Extended Data Fig. 8). Our RNA-seq datasets revealed distinct gene sets in *rca1*Δ and *efg1*Δ mutants implicated in cell wall organization^3,50^, adhesion^51^ and morphogenetic switching^17,52,53^ (Extended Data Fig. 7a, d, e, 8e). Hence, Rca1 and Efg1 may also affect immune recognition^3^ and *in vivo* fitness. To test colonization and fitness traits of *C. auris* strains, we used two primary mouse skin models (Fig. 4a, b)^54–56^. First, we tested whether the low CO_2_ concentration that *C. auris* encounters on the skin surface affects colonization *in vivo*^55,57^. Remarkably, after colonizing mouse back skin for 14 days, *rca1*Δ and *efg1*Δ mutants showed dramatically reduced fungal burdens when compared to the WT (Fig. 4a, Extended Data Fig. 10a). Similarly, striking effects were noted when *C. auris* colonized a primary native *ex vivo* human skin model as analysed by scanning electron microscopy (SEM) and CFU burdens (Fig. 4c, d). Most remarkably, fungal burdens were dramatically reduced by around 10-100-fold for both *rca1*Δ and *efg1*Δ mutants when compared to the WT (Fig. 4d). Subsequently, we investigated whether there is an impact on fitness *in vivo* when *C. auris* reaches deeper skin compartments where CO_2_ concentrations can reach up to 5.5%. Fungal cells were injected intradermally into mouse back skin. On day 3 post-infection, skin biopsies were digested and homogenized to quantify fungal burdens, unequivocally revealing fitness deficiencies within deeper skin compartments (Fig. 4b). One possible explanation could be that the marked cell surface alterations in *rca1*Δ and *efg1*Δ mutants render them more susceptible to phagocytosis and killing by the host immune defence (Extended Data Fig. 10b, Fig. 4b).

**Fig. 4.**
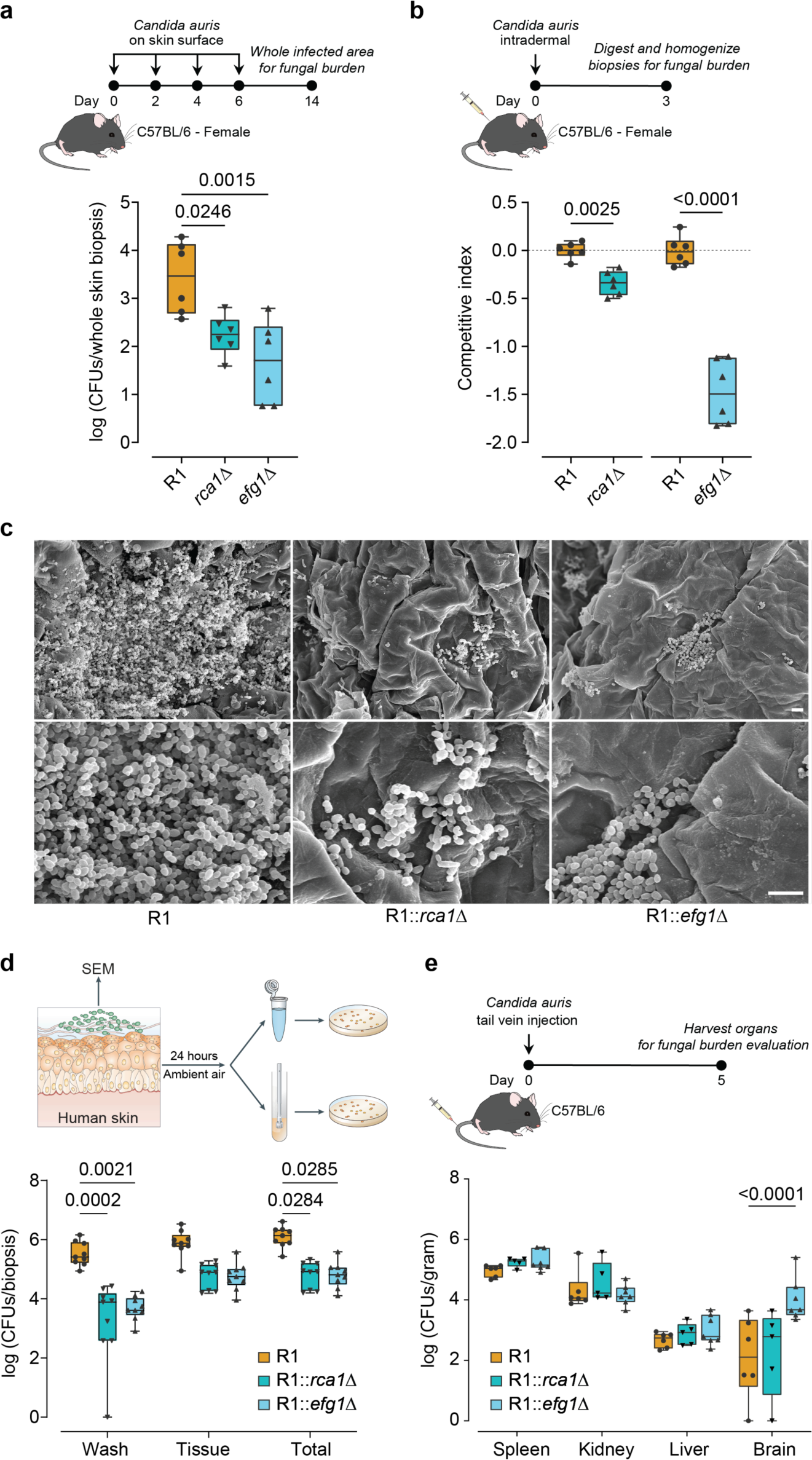
│ Rca1 and Efg1 are required for skin colonization and fungal fitness. **a**, Fungal burden in the skin biopsies after 14 days of colonization on back skin (n=6). **b**, Competitive fitness assays compared mutants and the WT strain following intradermal infection for three days (n=6). **c**, Representative images of scanning electron microscope (SEM) pictures of human skin surface topically infected with *C. auris* for 24 hours. Scale bar: 10µm. **d**, 3µl of fungal suspension were applied on human skin biopsies. After 24 hours, biopsies were washed in PBS and CFUs burdens were quantified in both washing supernatants and skin tissue samples, with the total representing the sum of burdens in both fractions. P values were calculated from average burdens of each donor. **e**, Fungal burdens in organs were quantified at day five post tail-vein fungal infections (n=5-6 female mice; see Extended Data Fig. 10c for male mice). P values were obtained by two-tailed unpaired t-test **(b)** or One-way ANOVA **(a)** or Two-way ANOVA **(d, e)** with Tukey’s multiple comparisons tests.

Due to the relatively thin nature of mouse skin compared to human skin, the dermal layers may not provide adequate CO_2_ levels to facilitate fitness recovery for *rca1*Δ. Consequently, we evaluated the performance of these strains in systemic infections and examined how fitness is sustained in deep organs where approximately 5.5% CO_2_ predominates. Unexpectedly, lateral tail vein injection of 10^8^ cells led to 100% mortality in mice infected with *efg1*Δ (n=3). We attribute this outcome to the severe flocculation trait of the *efg1*Δ mutant (Extended Data Fig. 8d) that led to cell aggregation and subsequent blockage of blood capillaries. Hence, we administered 2×10^6^ fungal cells into the lateral tail vein, followed by CFU quantification in different organs. Remarkably, with exception of the brain, we observed no significant differences in fungal burden between WT and *rca1*Δ or *efg1*Δ mutants in the spleen, kidney and liver in both female and male mice (Fig. 4e, Extended Data Fig. 10c).

These data demonstrate that CO_2_ levels in internal organs are essential and adequate to rectify *in vivo* fitness deficits caused by a dysfunctional CSP. Taken together, our results unveil a novel previously unrecognized critical role of the CO_2_-sensing pathway in governing both antifungal resistance and skin tropism of *C. auris*. The regulators Rca1 and Efg1 respond to tissue CO_2_ levels to control the crucial effector component Nce103. Therefore, Rca1 and Efg1 not only act as master regulators (Extended Data Fig. 7b, c, 11) of diverse metabolic processes and cell surface functions but also govern skin colonization and virulence of *C. auris*.

## 3. Discussion

The ease of person-person transmission constitutes a major challenge for proper clinical management of *C. auris* infections. Moreover, the pronounced skin tropism and inherent as well as acquired frequent pan-antifungal resistance traits thus prompted the WHO to declare *C. auris* as one of the top three fungal priority pathogens for drug development. Here, we show that a carbonic sensing pathway (CSP) is critical to establish growth of *C. auris* on the skin. Debilitating the CSP eradicates fungal fitness and thus virulence. Moreover, the CSP also controls resistance to the drug AMB. We identify the transcription factors Rca1 and Efg1 as master regulators of the Nce103 carbonic anhydrase, the key enzyme of the CSP converting CO_2_ into bicarbonate, to sustain energy metabolism when colonizing and growing on skin tissues in ambient air.

### 3.1. Exploiting a CSP to control AMB^R^ trait is unique in the *C. auris* skin pathogen

The *C. auris* Nce103 beta carbonic anhydrase is highly conserved in fungal species (Extended Data Fig. 4). This enzyme controls HCO_3_^-^ homeostasis and thus affects the downstream energy metabolism. Since Nce103 also operates in the cAMP/PKA pathway, it affects morphogenetic changes, drug resistance and virulence as reported before for many fungal pathogen species^30,31,31,34,37,58–61^. Although the architecture of the regulatory networks use conserved components, they differ in distinct fungal species concerning their specific roles in pathophysiology^36^. In *C. albicans*, deletion of *RCA1* slightly increases 5FC susceptibility, and enhances resistance to echinocandin and azoles^37^. This mirrors overexpression of the carbonic anhydrase Can2 in *C. neoformans,* leading to polyene and azole hypersensitivity^32^. Remarkably, we show here that the CSP exploits Nce103 to augment AMB^R^ in *C. auris* isolates from different clades (Fig. 2d, Extended Data Fig. 3d). Ablation of *RCA1* or *EFG1* re-sensitizes *C. auris* to AMB as it leads to a substantial 2-4-fold decrease in AMB^R^, which is consistent with a 2-4-fold increase when overexpressing *NCE103* (Fig. 2f). Of note, the CSP-mediated control of AMB^R^ traits appears as a specific trait of *C. auris* not observed in other fungal pathogens (Fig. 2e). Indeed, MIC assays on other skin-resident fungal pathogens such as Malassezia and Dermatophyte (Extended Data Fig 12a, b) show no changes in the AMB response between 5.5% CO_2_ and ambient air. These differences may relate to distinct nutrient requirements of Malassezia and Dermatophytes, which seem to scavenge host lipids and keratin as main energy sources^62,63^.

The obvious impact of CO_2_ levels on AMB^R^ phenotypes of *C. auris* may also have serious ramifications for clinical MIC testing. Hence, our data imply a critical reassessment of standard clinical AMB susceptibility testing protocols for *C. auris*^64^, and perhaps incorporating CO_2_ supplementation in the MIC-testing. Most importantly, the pivotal role of Nce103 in the control of *C. auris* AMB susceptibility suggests that Nce103 inhibitors could be promising for topical applications to clear *C. auris* colonizing skin surfaces. This would also minimize skin-to-skin transmission and facilitate clinical management of colonized patients. AMB is fungicidal and the drug has severe adverse effects limiting its long-term use^46^. AMB^R^ has been rare for decades, most likely due to its fungicidal action and severe fitness trade-offs^65^. However, *C. auris* can rapidly acquire clinical resistance, with up to 50% of clinical isolates showing untreatable AMB^R^ ^22^. While drug resistance traits often come at fitness trade-offs, *C. auris* can acquire AMB^R^ through mutations in the ergosterol biosynthesis pathway^15^. The fitness cost can be overcome by a frameshift variation in *CDC25*, encoding an activator of Ras/adenylyl cyclase pathway^15^. Of note, AMB^R^ without significant fitness trade-offs constitutes a severe clinical challenge for *C. auris* treatment, as AMB^R^ at physiological CO_2_ levels in combination with inherent azole and echinocandin resistant traits favour pan-antifungal resistant pathogens, which are indeed increasing in prevalence^5,66^.

### 3.2. The CSP in *C. auris* is essential for skin colonization

Our data shows that the CSP is necessary for *C. auris* skin colonization in mammals. The human skin surface typically holds higher salt levels creating osmostress but poor nutrient availability^67^. Indeed, while *rca1*Δ exhibits only minor sensitivities to osmostress, both *efg1*Δ mutants and strain S, which lacks a functional Rca1 protein, show clear osmosensitivity phenotypes (Extended Data Fig. 6a). Hence, it is tempting to speculate that salinity stress *C. auris* encounters on the skin surface also engages Efg1 to cope with osmotic fluctuations (Extended Data Fig. 6a), in addition to the hallmark Hog1-dependent osmostress responses^68,69^. Moreover, all skin-tropic microbes must exploit strategies to deal with nutrient deficiencies and persistent stress. In the case of *C. auris*, it can utilize CO_2_ as carbon to ensure energy homeostasis and fitness of *C. auris* in the deprived nutrient environment of the skin surface. Similar to other microorganisms, *C. auris* R1 exhibits the same capacity to generate biomass in YP with/without glucose medium (Extended Data Fig. 9e). Hence, secreted aspartic proteases (SAPs) and the gluconeogenesis pathway may play supportive roles in its nutrient acquisition strategies on the skin. However, Rca1 and Efg1 enhance growth in medium lacking glucose, thereby emphasizing the essential role of CSP for *C. auris* fitness (Extended Data Fig. 9e). Furthermore, the deletion of *RCA1* or *EFG1* leads to the upregulation of various iron metabolism genes, including *SIT1, FET33, FTR1,* and *AIM14* (Supplementary Table 6). This suggests a putative role for the CSP in iron scavenging of *C. auris* on the skin, where typically lower iron concentrations prevail when compared to deeper skin layers or other body tissues^70^. Indeed, iron-limitation is a serious problem for fungal growth, and thus fungal pathogens often scavenge essential trace metals from host cells or tissues^71^. Although the skin surface is a nutrient-limited niche, the pronounced skin tropism of *C. auris* has several explanations. First, the skin’s immune defense on the surface is less active and unable to provide full protection, thus making the niche less hostile for pathogens. Second, systemic antifungal drugs do not reach pharmacological active concentrations on the skin surface. Finally, the metabolic flexibility of *C. auris* may offer additional benefits, since the pathogen would be able to utilize metabolites released by skin microbiota as alternative nitrogen or carbon sources. Hence, the CSP may also maintain *C. auris* fitness on the skin niche through beneficial interactions with other commensal microbes, including *Proteus mirabilis, Pseudomonas aeruginosa, Klebsiella pneumoniae, Providencia stuartii* and *Morganella morganii*^10,72^. For example, *C. auris* could benefit from those skin-biome bacteria that use urease to generate CO_2_ from urea, which is naturally present on human skin and in sweat^67^. The competition for nutrients in such shared niches also has the potential to shape the dynamics and equilibria of microbiome compositions. Thus, we tested how CO_2_ affects *Malassezia* spp., potential competitors of *C. auris* residing on human skin tissues^10^. Remarkably, 5.5% CO_2_ enhances the growth of *M. furfur* while inhibiting *M. sympodialis* (Extended Data Fig. 12a). This observation indicates that different fungal species exploit distinct strategies for nutrient acquisition when sharing the same niche. This is biologically relevant, since fitness may be compromised when microbes rely on similar nutrient acquisition strategies within the same niche. Therefore, we believe that CO_2_-driven nutrient acquisition by *C. auris* plays critical roles for colonizing and growing on native skin, but also reduces competition with other skin microbes (Extended Data Fig. 13).

### 3.3. A dual function of the CSP in the control of energy metabolism and antifungal resistance

Initially, we hypothesized that the CSP regulates AMB^R^ traits by affecting energy metabolism, as glucose could restore the growth of strains S and *rca1*Δ (Fig. 3e). Unexpectedly, the presence of 2% glucose in culture medium does not alter the MIC for AMB when compared to RPMI or YP alone (Extended Data Fig. 3g). Hence, the control of energy metabolism and the development of AMB^R^ traits may be due to independent functions regulated by CSP. On the other hand, supplementing 5.5% CO_2_ *in vitro* efficiently restored AMB^R^ in both *rca1*Δ and *efg1*Δ mutants. Our RNA-seq data show that both Rca1 and Efg1 control expression of *NCE103*, which may in part explain why CO_2_ supplementation restores AMB^R^ in *efg1*Δ. However, it remains unclear why CO_2_ fails to restore apparent fitness defects of *efg1*Δ mutants (Fig. 3e, Extended Data Fig. 9b). We speculate that morphogenesis may play a role, as Efg1 is also a hallmark regulator of morphotype variations in relevant *Candida* spp., which also engages the PKA signalling pathway^33,59^. Thus, HCO_3_^-^ may directly activate the cAMP/PKA cascade through Cyr1, but we cannot rule out the possibility that other factors are acting on the PKA complex Tpk1-Tpk2-Bcy1. Surprisingly, deletion of *CYR1* in *C. auris* B8441 reduces AMB susceptibility, but this effect is not observed in *tpk1*Δ, *tpk2*Δ, or *tpk1*Δ*tpk2*Δ mutants^18^, suggesting complex genetic interactions. Hence, additional genetic studies are required to verify whether an alternative *C. auris* pathway can associate CO_2_-sensing with Efg1 activity, or if CO_2_ can directly control Efg1 downstream function in the context of AMB resistance.

Our RNA-seq data uncover distinct transcriptional profiles subject to control by Rca1 and Efg1 (Extended Data Fig. 6d). Thus, we believe that Rca1 is acting upstream in the CSP to control Nce103, but engages Efg1 depending on the metabolic state, the presence of immune defence or the need to undergo morphogenetic changes within deeper skin compartments^56^. Additional prominent functions influenced by CSP include regulation of the cell wall architecture, membrane lipid alterations as well as metabolic pathways (Extended Data Fig. 8, 11). Moreover, *rca1*Δ and *efg1*Δ strains carry a substantial enrichment of RNA-processing and ribosomal biogenesis genes (Extended Data Fig. 14a, b). Although enriched genes are distinct between *rca1*Δ and *efg1*Δ mutants (Extended Data Fig. 14c), Rca1 and Efg1 could still control post-transcriptional as well as translational regulation on specific subsets of target genes.

## 4. Conclusion

In this study, we discover hitherto unrecognized functions of the CSP in controlling antifungal drug resistance and skin colonization of *C. auris*. We demonstrate that the carbonic anhydrase Nce103 can fix ambient CO_2_ to support energy metabolism and to sustain fungal growth when colonizing skin tissues. Further, the CSP engages downstream Efg1 to maintain fungal homeostasis and fitness of *C. auris* under host defence on and within skin tissues. Importantly, Rca1 and Efg1 cooperate to regulate *NCE103* encoding carbonic anhydrase. The data support the notion that Nce103 is a suitable antifungal target in therapeutic approaches to control or prevent skin colonization by *C. auris* through topical applications. Taken together, this work provides comprehensive mechanistic insights into how a CSP contributes to fungal virulence, fitness and polyene resistance traits in *C. auris*.

## 5. Acknowledgements

We thank Andriy Petryshyn, Semiye Volkan and all lab members for technical support and experimental advice. Moreover, we thank Neeraj Chauhan for providing clinical isolates; Axel A. Brakhage for *Trichophyton benhamiae* (WT and *stuA* mutants); Salomé LeibundGut-Landmann and Joseph Heitman for advice on Malassezia experiments; Clarissa Nobile for mentoring as a PhD Thesis Committee member; Rounik Mazumdar for supporting proteomics experiment; Ashton Trey Belew for consulting on differential expression analysis; Phuc-Loi Luu and VnPathoInformatics for guide in bioinformatics. We are grateful to all volunteers who donated skin tissues for this study. This work was supported grants from the Austrian Science Fund FWF to K.K. (ChromFunVir; P-32582-B08, Candidomics P-33425, BacFun P-34152 and FWF-SFB70-08). A.E.-B. work received funds from the Austrian Science Fund (FWF; P 31485-B30) and the LEO Foundation (LF-OC-23-001332). P.C.-T. was supported by a Student Fellowship, *Ernst Mach Grant*, from ASEA-UNINET network; the ESCMID Research Grant 2023 from the European Society of Clinical Microbiology and Infectious Diseases; H.A. and P.C.-T were supported by the FWF-funded PhD training program (*TissueHome* – FWF-DOC32-B28).

## 6. Author contributions

P.C.T. and K.K. formulated the study, designed experiments, analysed and interpreted data, wrote the manuscript. K.K: PhD supervision and funding management. P.C.-T. conducted most experiments, data analysis, as well as visualization. A.E.-B. coordinated the arrangement of skin samples, provided methodology, and contributed to data interpretation. S.S. and D.C. facilitated human skin experiments. S.S. and D.M. performed SEM. P.P. aided systemic infections and macrophage interaction experiments in mice. T.B. helped with RNA experiments. H.A. helped with the mouse back skin models. W.C. and M. H. assisted with proteomics and data analysis. A.K. helped with fungal gene deletion. S.J. helped with cloning and data analysis. N.K. performed computational structural modelling and docking. L.M.Z., M.L. and C.M. performed sterol lipid analysis and helped with antifungal susceptibility testing. G.I. supported Malassezia experiments. A.C. provided clinical strain collections and helped with strain typing. A.E.-B. conducted manuscript review and editing.

## 8. Methods

### 8.1. Ethics statement

Animal experiments adhered to ethical approval from the ethics committee of the Medical University of Vienna and the Federal Ministry of Science and Research, Vienna, Austria (BMBWF-66.009/0436-V/3b/2019). Adult wild-type C57BL/B6 mice were housed in specific pathogen-free conditions at the Max Perutz Labs Vienna, Vienna BioCenter Campus. Abdominal human skin samples were obtained from anonymous healthy adult female donors (28-59 years) with approved consent following the Declaration of Helsinki and ethics committee approvals of the Medical University of Vienna (ECS 1969/2021).

### 8.2. Fungal growth conditions, media formulations and dose-response assays

A list of fungal strains used in this study is provided in Supplementary Table 1. Clinical *C. auris* isolates were obtained from CDC & FDA Antimicrobial Resistance (AR), Indian National Culture collection of Pathogenic Fungi (a kind gift from Neeraj Chauhan and Anuradha Chowdhary). All isolates were preserved at –80 °C in 25% glycerol, and freshly streaked on YPD agar for experiments. *C. auris* was routinely grown YPD medium at 30 or 37 °C, 200 rpm. For growth curve experiments, we used amino acid drop-out media, and minimal medium or yeast-peptone (YP) with different carbon sources. All medium compositions were provided in Supplementary Table 2. All chemical reagent vendors and identifiers were provided in Supplementary Table 3.

Growth curves were performed for different culture media formulations as described in Supplementary Table 2. Fungal cells from an overnight culture in YPD at 30 °C, 200 rpm were washed 3 times with distilled water (dH_2_O), followed by measuring OD_600nm_. Cell suspensions were adjusted to 0.2 OD_600nm_ in dH_2_O. Aliquots of 100 µl of testing media (two times concentration) were dispensed into non-treated 96-well plates (Starlab). Subsequently, 100 µl of fungal suspensions were added to each well. Plates were incubated in static incubators at 37 °C with/without 5.5% CO_2_. The OD_600nm_ was recorded every two hours within three days by Victor Nivo plate reader (PerkinElmer).

Dose-response assays were done adhering to the CLSI M27-A3^73^ protocol in 96-well plates with some minor modifications. Briefly, yeast cells from an overnight cultured were reinoculated into fresh YPD medium at initial 0.2 OD_600nm_. After a four-hour incubation, fungal cultures were adjusted to OD_600nm_ of 0.1 in distilled water. A 25 µl aliquot of fungal suspensions was diluted into 10 ml RPMI 1640 (Gibco) buffered with 35 g/l MOPS (AppliChem), pH 7 (RPMI). The 96-well plates containing 100 µl media with drug compounds were prepared by two-fold serial dilutions. The 100 µl of the fungal suspensions was aliquoted into each well. Negative controls lacked inoculum, whereas positive controls included inoculum without adding agents. Plates were incubated at proper conditions following experimental requirements. Optical density OD_600nm_ was recorded in Victor Nivo Microplate Reader. IC_50_ was calculated using the four-parameter log-logistic function (LL.4) from *drc* package^74^ in R. Drugs were prepared in 100x stock solution in DMSO or dH_2_O. Reagent stocks DMSO were used at a maximal a final concentration of 1% DMSO. 100% growth inhibitory concentration (MIC_100_) values were also collected visually after 48 hours. Similar protocols with some modifications were used for Malassezia, and filamentous fungi were described in Supplementary information (SI 1).

### 8.3. Proteomics

Several colonies of *C. auris* growing on YPD agar were picked and re-grown overnight in RPMI. Fungal suspensions were transferred to 50 ml fresh RPMI to reach OD_600nm_ of 0.1 in baffled flasks. After seven hours of incubation at 30 °C with agitation of 200 rpm, yeast cells were pelleted and washed three times in cold PBS (Sigma-Aldrich). Protein was extracted and digested before subjecting to LC/MS-MS analysis following the workflow presented in Supplementary Information 1 (SI1). Raw data were analysed with MaxQuant software (version 1.6.17.0)^75^ using the Uniprot *C. auris* reference proteome (version 2021.03). Output tables were further processed in R using Cassiopeia_LFQ (https://doi.org/10.5281/zenodo.5758974) before subjecting into differentially protein analysis with LIMMA package^76^.

### 8.4. RNA isolation, qPCR and RNA-sequencing

Fungal cultures were centrifuged, and cell pellets were rapidly frozen in liquid nitrogen. Dry cell pellets were then stored at –80 °C for later use. Total RNA was extracted using the TRIZOL method, followed by DNase I treatment procedure with protocol provided in SI1. RNA quality and purification was checked with Nanodrop and conventional PCR-based quantification of *ACT1* mRNA. For quantitative PCR, first-strand cDNA was synthesized from RNA with Reverse Transcription System Kit (Promega). Subsequently, 15 ng cDNA was utilized for qPCR amplification, employing the 2x Luna Universal master mix (NEB). For competitive assays, gDNA was used directly for qPCR. The data were analysed using the cloud-based system provided by Bio-Rad accessible at BR.io.

For RNA sequencing (RNA-seq) analysis, fungal cells grown overnight were diluted into 15 ml fresh YPD to reach OD_600nm_ of 0.1 in baffled flasks. Flasks were then incubated at 37 °C until reaching OD_600nm_ of 2.5. Fungal cells were collected by centrifugation, and RNA was isolated following protocol in SI1. RNA quality and integrity were evaluated with Bioanalyzer RNA 6000 Nanochip (Agilent Technologies). The library and sequencing were performed at the commercial Novogene Sequencing Facility (UK). Briefly, mRNA was enriched by poly-T oligo-attached magnetic beads followed double-stranded cDNA library preparation. The quality-controlled RNA libraries were pooled and sequenced with 150-bp paired-end reads on the Illumina NovaSeq 6000 platform.

RNA-seq bioinformatics data analysis used a workflow established before^19^ with some minor modification (https://github.com/kakulab/CSP2024). Briefly, quality of raw RNA-seq data was assessed by fastQC v0.11.9^77^. TrueSeq (Illumina) adapters were trimmed by cutadaptv2.8^78^ (settings –interleaved –q 30), followed by read mapping on *C. auris* B8441 genome assembly (Candida Genome Database version s01-m01-r22) using NextGenMap v0.5.5^79^ (settings –b –Q 30). rRNA loci were removed by BEDtools v2.29.1^80^ (settings: intersect –a –b –v). Then, Picard tools ^81^ from Broad Institute was used to remove duplicate reads (settings: MarkDuplicates REMOVE_SEQUENCING_DUPLICATES=true). HTSeq^82^ was used for read counting in the union mode (settings htseq-count –f bam –r pos –t gene –i ID). Genomic annotation of *C. auris* B8841^83^ version s01-m01-r22 was used. SAMtools v1.15.1^84^ was used to prepare coverage files for Integrative Genomics Viewer^40^ (IVG) visualization. Differentially gene expression (DEG) analysis was performed as pairwise comparisons using EdgeR v3.40.2^85^. The adjusted p value to control the false discovery (FDR) computed from p values using the Benjamini-Hochberg method. Normalized read counts were calculated by cpm function in edgeR which input into prcomp function in *stats* v3.4.1 in R for principal component analysis (PCA).

### 8.5. Generation of *C. auris* deletion mutants

*C. auris* deletion mutants were generated by gene replacement as described before^86^ using a gene-specific a deletion cassette with a dominant marker constructed using the three-way stitching PCR method^86^. Briefly, approximately 500 bp upstream and downstream flanking regions of the target gene were amplified from genomic DNA of *C. auris* strains. The *NAT1* selection marker flanked by the *TEP* promoter and terminator was amplified from plasmid pTS50^86,87^. DNA fragments were purified by GeneJET gel extraction kit (ThermoScientific), followed by a stitching PCR to obtain gene deletion cassettes for genomic targeting. PCR products were transformed into *C. auris* by electroporation protocol described before^86,88^. Colony PCR with OneTaq 2X master mix (NEB) were used to verify loss of target gene and correct cassette integration^86^. For overexpression of *NCE103*, we placed the *NCE103* locus under the control of the *ENO1* promoter from *C. auris*. The cassette containing *urNEUT1*-*pENO1-NCE103-pTEP1-NAT1-drNEUT1* was cloned into a long intergenic region (*NEUT1*) on chromosome 1 located between B9J08_000423 and B9J08_000424. All oligonucleotide primers and plasmids are listed in Supplementary Table 4 and 5.

### 8.6. Homology searches, multiple sequence alignments and phylogenetic tree construction

DNA sequences retrieved from CGD^83^ were subjected to *blastn* from NCBI. Homology was searched on nucleotide collection (nr/nt) and whole-genome shotgun contigs (wgs) for *C. auris* (taxid:498019). Searching was done with the coding sequences for *RCA1, EFG1, NCE103*. Detailed results can be found in Source data.

Protein sequences of *Candida* spp. were retrieved from CGD^83^. *Blastp* searches were performed on *C. auris* proteins to find best hits in target species. A reciprocal search against *C. auris* genome was conducted on *bastp* to identify orthologous genes. All sequences were put into *msa* package version 1.32 for multiple sequence alignments^89^. Phylogenetic trees were constructed with neighbour-joining method using *nj* function in ape package version 5.7^90^, followed by a bootstrap analysis with *boot.phylo*^90^. The *ggtree* package^91^ were used for phylogenetic tree visualization.

### 8.7. Structure prediction and molecular docking analysis

Beta carbonic anhydrase structures of *C. albicans* (PDB: 6GWU and UniprotKB: Q5AJ71) and *Coccomyxa* (PDB: 3UCJ) were retrieved from the PDB and UniprotKB repositories. Three computational homology models of *C. auris* Nce103 were generated using the SWISS-MODEL tool^92^ using the coordinates for Q5AJ71 (model 1), 6GWU (model 2) and 3UCJ (model 3). The molecular visualization and superimposed models were analysed by PyMOL version 2.5.2. Molecular docking was performed using CB-Dock (Cavity-detection guided Blind Docking)^93^. Inhibitor structures for acetazolamide and ethoxzolamide structures were retrieved from the PubChem database.

### 8.8. Assays for intracellular reactive oxygen species

Fungal cells from fresh overnight cultures were reinoculated into fresh YPD medium to reach 0.2 OD_600nm_ for four hours at 30 °C. Fungal cells were then washed twice and adjusted to 4×10^7^ cells/ml in PBS. Serial two-fold dilutions of AMB from 0.5-16 µg/ml were prepared in PBS and 100 µl aliquots of each concentration were dispensed into a 96-well plate. Subsequently, 100 µl of fungal suspensions were distributed into wells then incubated for one and a half hours at 37 °C. Dihydroethidium (DHE) was added at a final concentration of 25 µM and incubated for additional 30 minutes (min). Finally, fungal cells were washed twice, and ROS was measured by flow cytometry BD Fortessa (PE-TexasRed, 10,000 events). Single cell populations were exported using the FlowJo v10.8 software and visualized with ggplot2^94^. Mean fluorescent intensity (MFI) of PE-TexasRed signals were also exported in Extended Data Fig. 6.

### 8.9. Fluorescein diacetate (FDA) uptake

Fungal cells from overnight cultures were grown to the exponential growth phase in YPD starting from an initial OD_600nm_ of 0.2. Cells were washed twice in FDA buffer (50 mM HEPES, pH 7.0 and 0.5 mM 2-deoxy-D-glucose; Sigma-Aldrich) followed by quantification in a CASY cell counter. Fungal cell suspensions were adjusted to 5×10^6^ cells/ml in FDA buffer, before adding fluorescein diacetate (Sigma-Aldrich) at a final concentration of 50 nM. Aliquots of 200 µl of cell suspensions with/without FDA were immediately transferred to a 96-well optical-bottom plate (ThermoScientific). The kinetics of FDA uptake was followed by recording fluorescence every 5 min for 35 cycles at 37 °C in a Victor Nivo Microplate Reader with an excitation/emission setting of 485/535nm.

### 8.10. Mouse and human skin models, colonization assays and scanning electron microscopy

Animals used for experiments were WT C57BL/6 mice aged 8 to 14 weeks at the start of experiments. Female mice were employed for skin colonization experiments, while both female and male mice were used for systemic infections via the lateral tail vein. For skin colonization assays, mice were anesthetized with ketamine-xylazine (100 mg ketamine/kg body weight and 4 mg xylazine/kg body weight), followed by hair removal from back skin using an electric shaver to generate an area ∼ 9 cm^2^. A total of four applications were conducted on the area using suspensions containing 2×10^8^ fungal cells (WT and mutant strains) in 100 µl PBS, with fungal cells applied every two days. Mice were sacrificed on day 14 by cervical dislocation, and fungal cells on the skin surface were collected by a cotton swab moistened with 80 µl PBS, then immediately plated on YPD agar containing ampicillin, tetracycline, and chloramphenicol (YPD-CAT). Skin at infection area was excised and dissociated in 500 µl of enzyme solution containing 1 mg/ml Collagenase Type II (Gibco), 1 mg/ml DNase I (Roche), incubated at 37 °C, 5.5% CO_2_. After one and a half hours, samples were homogenized with a homogenizer (IKA 3386000) for 30 seconds, followed by plating onto YPD-CAT agar plates to quantify CFUs. For competitive assays, 2×10^8^ cells of both WT and mutant strains were mixed in 100 µl PBS and applied to the back skin surface. Fungal cells were again recovered as before and subjected to gDNA extraction and qPCR.

For intradermal infection, an equal cellular mixture of 5×10^6^ WT and mutant cells in 15 µl PBS were injected into the shaved back skin after anesthesia. On day three post infection, mice were euthanized by cervical dislocation, and 8 mm diameter skin biopsies were obtained of infected sites using a KAI Biopsy Punch tool (Heintel – 29045435). Skin biopsies were homogenized as described above, followed by plating on YPD-CAT agar. Plates were incubated at 37 °C with 5.5% CO_2_ for 48 hours. Fungal cells were collected from plates for gDNA extraction and qPCR to identify the ratio of mutants and WT in each sample using the primers for *RCA1, EFG1* and *NAT1.* Fungal mixtures used for infection were plated to control the actual infection dose. Relative gDNA abundance of WT (primers for *RCA1 or EFG1*) and mutants (primers for *NAT1*) were calculated based on *ACT1* as a control gene. The relative gDNA abundance was adjusted with abundance ratio of input samples before calculating competitive values as log_2_ (relative abundance values to WT). For systemic infections, we injected 2×10^6^ fungal cells in 100 µl PBS per 21.5-gram mouse body weight via the lateral tail vein injected of WT mice^95^. On day 5 post infection, mice were sacrificed by cervical dislocation, and CFUs in internal organs (spleen, kidney, liver, brain) were quantified by plating^95^.

Native adult human skin samples (female donors, age range 28-59 years) were obtained within one to two hours post plastic surgery procedures^56^. Initially, the skin underwent a cleansing process using PBS, before being punctured with an eight mm KAI Biopsy Punch tool (Heintel – 29045435). Skin biopsies were then placed onto a 12-well plate containing Dulbecco’s Modified Eagle’s Medium (DMEM), supplemented with 10% fetal bovine serum (FBS) and 1% penicillin-streptomycin. Fungal cells from exponential growth phase cultures were washed twice with PBS, and subsequently adjusted 10^5^ cells/ml cell densities using a CASY cell counter (Roche). Three µl of fungal cell suspensions were applied topically to the center of the biopsies, with PBS serving as non-infected control and incubated in tissue culture plates at 30°C under ambient air. After 24 hours, biopsies were collected and washed in 1 ml of PBS by vortex-mixing for 10 seconds. Subsequently, biopsies were sectioned into four pieces and subjected to digestion with a 500 µl enzyme solution containing 1 mg/ml Collagenase Type II (Gibco), 1 mg/ml DNase I (Roche), and incubated for three hours at 37°C, 5.5% CO_2_. Next, 500 µl PBS was added and biopsies were homogenized with a mechanical tissue homogenizer (IKA 3386000) for 30 seconds. Washing suspension and homogenized biopsies were diluted and plated onto YPD-CAT agar for fungal burden assessment.

The preparation of human skin biopsies for SEM analysis was performed essentially as described previously^96^. In brief, biopsies were harvested after 24 hours and fixed in Karnovsky’s fixative (2% paraformaldehyde, 2,5% glutaraldehyde in 0.1 M phosphate puffer pH 7.4 from Morphisto^®^, Germany) for at least 24 hours. Samples were then washed two times in 0.1 M phosphate buffer at 4 °C (pH 7.3) for 2 min each. All samples were dehydrated in a graded ethanol series (50%, 70%, 80%, 90%, 95% and 100%) for 20 min each. Samples were immersed for 30 min in pure hexamethyldisilazane (HMDS, Sigma-Aldrich) followed by air-drying. For SEM analyses, samples were sputter-coated with gold (Fisons Instruments Polaron Sputter Coater – SC7610) and examined with a scanning electron microscope (JSM 6310, Jeol Ltd.^®^, Japan) at an acceleration voltage of 15 kV.

### 8.11. Statistical methods

Statistical analyses were conducted with GraphPad Prism version 9.0 and *rstatix* R package version 0.7.2^97^. Unless otherwise specified, data represent means ± standard deviation from at least three biological replicates. For mouse experiments, at least 5 mice were used for each group. For human skin experiment, data were collected from at least 3 independent human donors. Significance was determined using Student’s t-test or ANOVA, followed by Benjamini-Hochberg or Tukey’s or Dunnett’s post hoc tests for multiple comparisons. Detailed methods for Data integration and Extended Data Figures were described in SI1.

### 8.12. Data availability

The proteomics data have been deposited to the ProteomeXchange Consortium via the PRIDE partner repository^98^ with the dataset identifier PXD048342. RNA-seq data are available from the Gene Expression Omnibus (GEO) database with the accession number GSE253332. The code for RNA-seq analysis workflow was deposited at https://github.com/kakulab/CSP2024.

---

**Extended Data Fig. 1.**
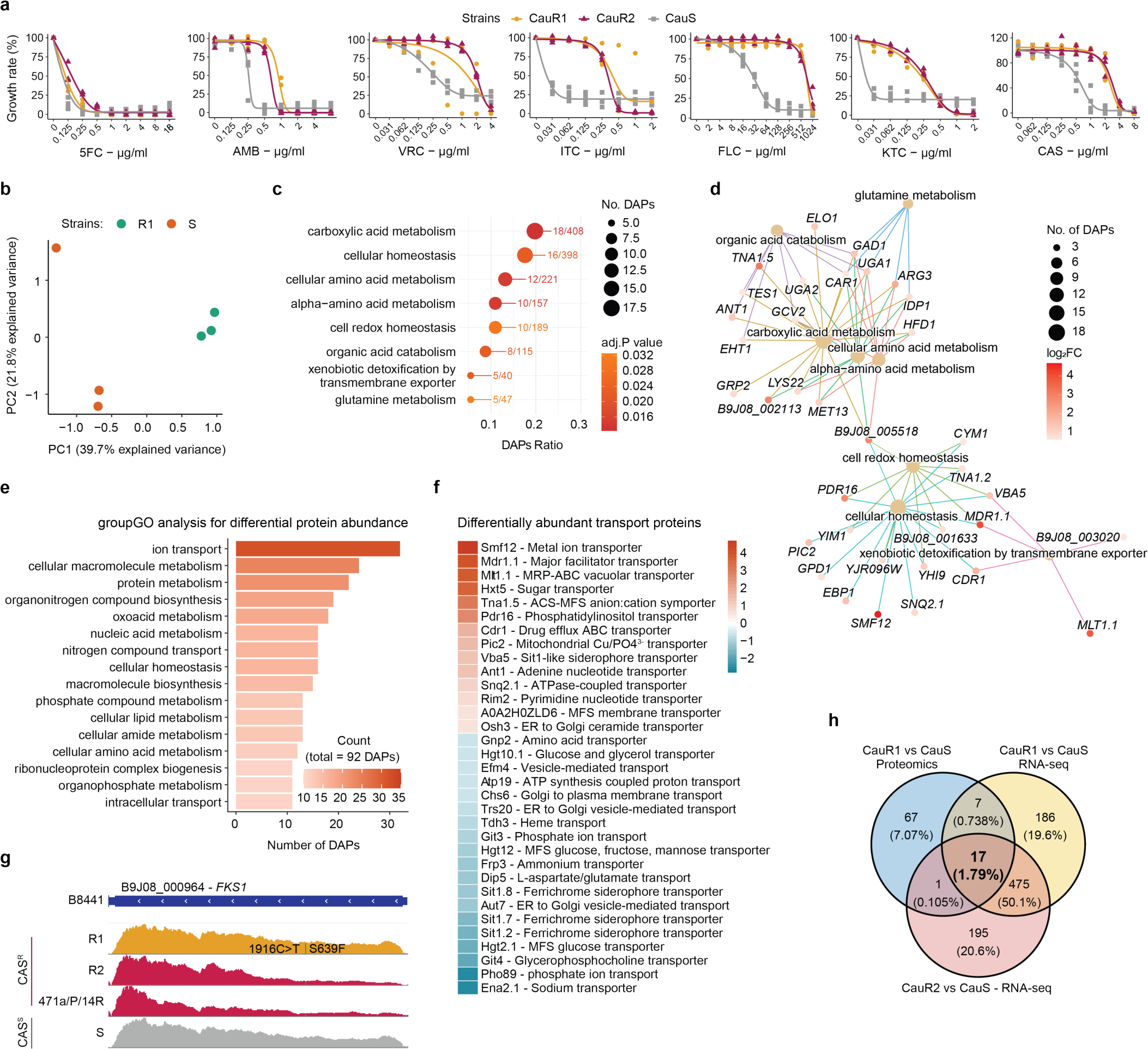
│ Integration of multi-omics datasets reveals potential drug resistance mechanisms. **a**, Dose-response data of resistant and sensitive strains for antifungal drugs in Fig. 1a. (amphotericin B – AMB, caspofungin – CAS, 5-fluorcytosine – 5FC, ketoconazole – KTC, itraconazole – ITC, fluconazole – FLC, voriconazole – VRC). **b**, PCA analysis of proteomics data from resistant and sensitive strains with 3 biological replicates for each strain. **c-d**, GO-enrichment analysis for differentially abundant proteins (DAPs). **e**, groupGO analysis, using clusterProfiler package reveals proteins involved in ion transport (32/92 DAPs) enhanced in resistant strains. **f**, Comparative proteomics reveals high abundance transporters between resistant and sensitive strains. **g**, IGV reveal a hotspot S639F variant (green) in Fks1 governing caspofungin resistance in R1, R2, 471/P/14Rj (Jenull et al. 2022). **h**, Integrative Venn diagram showing overlap of highly abundant proteins (see Fig. 1d) and highly expressed genes between R and S *C. auris* strains. **(c-f, h)** Data were exported from differential expression analysis of proteomics and RNA-seq datasets using a cut-off ± 0.58 for log_2_FC and adjusted p-value < 0.05 (calculated by Benjamini-Hochberg).

**Extended Data Fig. 2.**
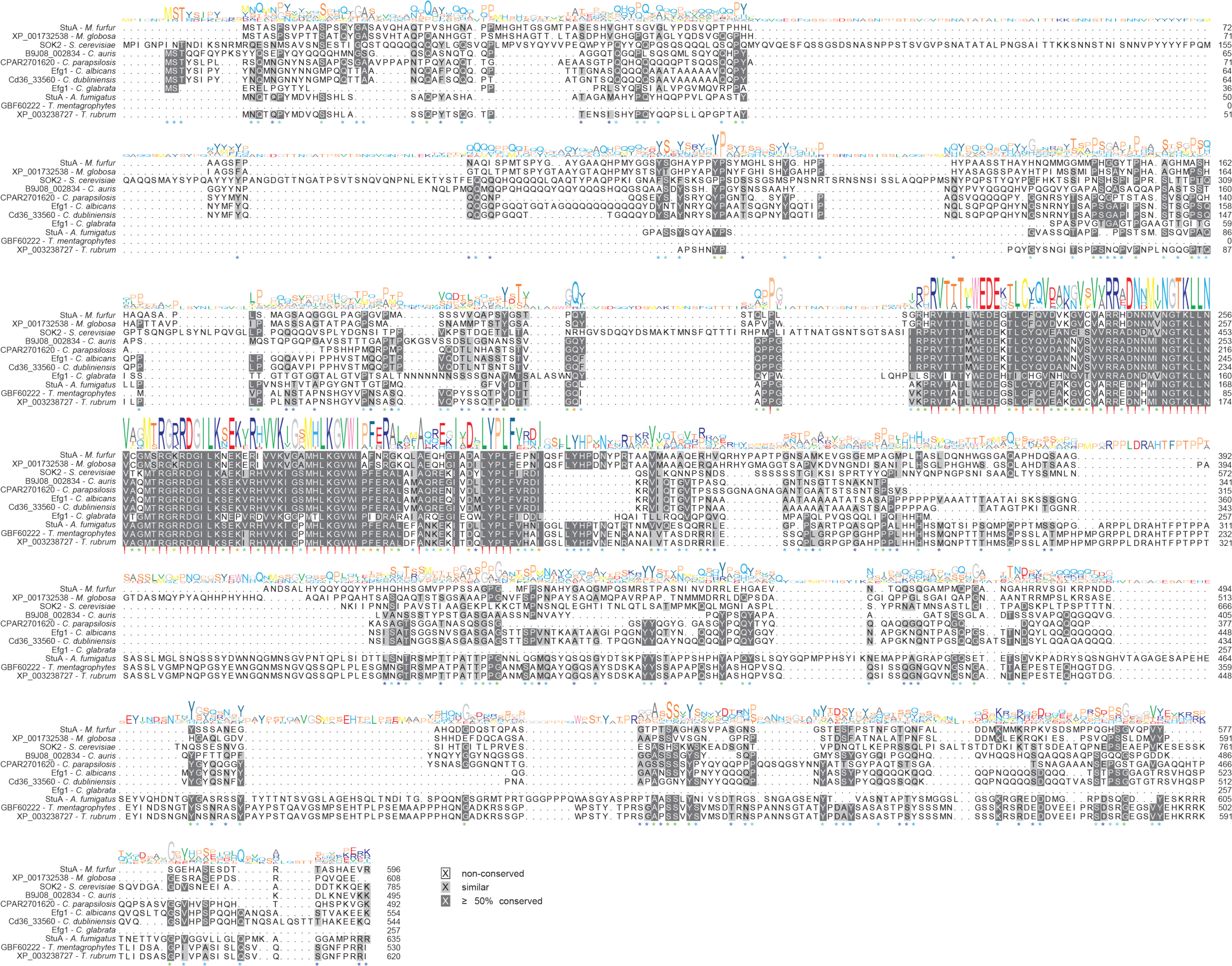
│ Multiple primary sequence alignments of Efg1 homologues across fungal pathogen species. Protein sequences were subjected to NCBI *blastp* searches on *C. auris* proteins sequences. The best hits were reciprocally searched against *C. auris,* all of which confirm that the open reading frame B9J08_002834 encodes the transcriptional regulator Efg1.

**Extended Data Fig. 3.**
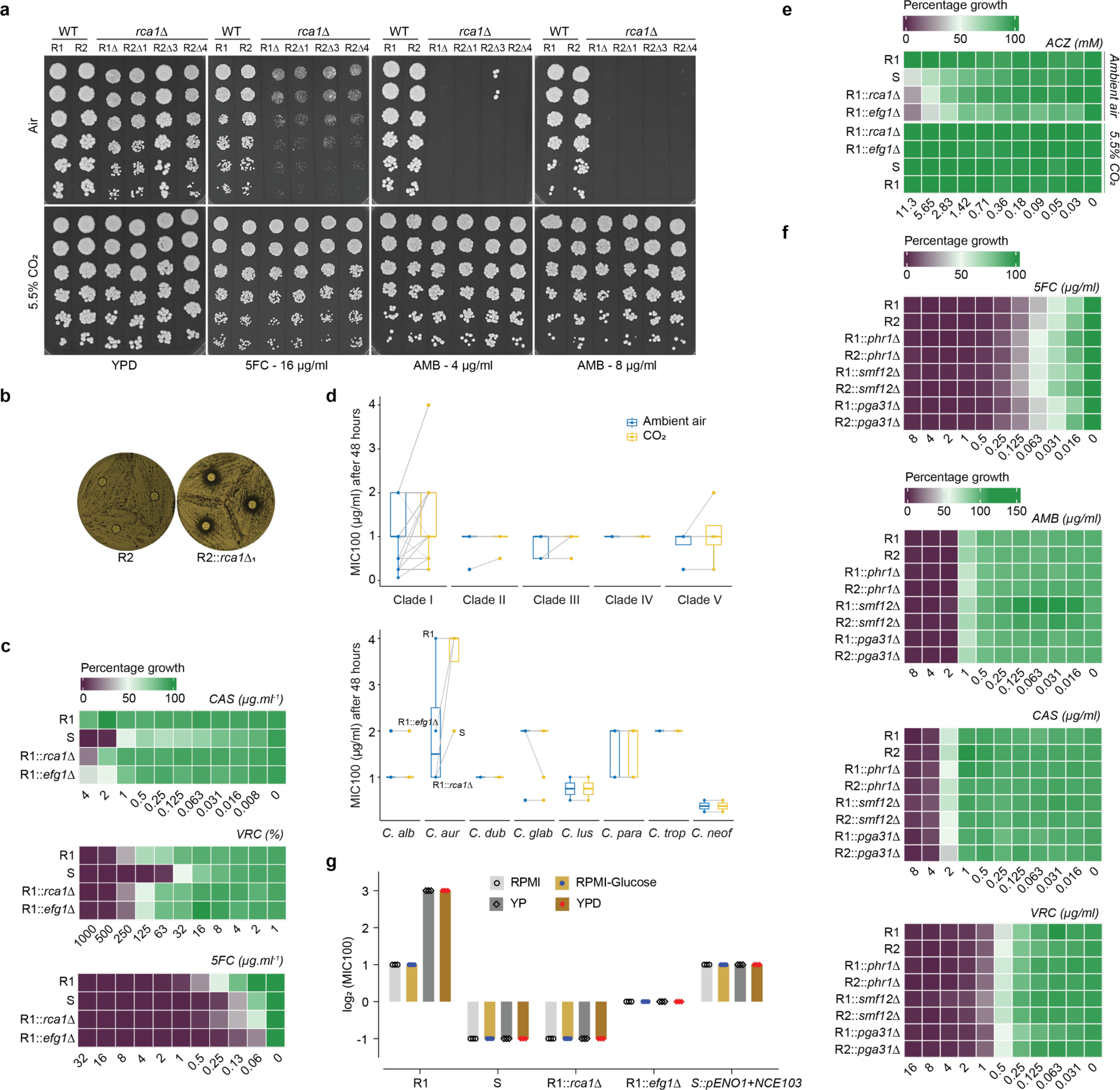
│ The CSP controls drug resistance in *C. auris*. **a**, Suspensions of 5×10^7^ fungal cells/ml were five-fold serially diluted. Aliquots of 1µl of cell suspensions were spotted onto YPD agar or YPD agar containing antifungal drugs. Pictures were taken after growing cells for 48 hours at 37 °C in ambient air or 5.5% CO_2_. **b**, Fungal cells were adjusted to OD_600nm_ ∼ 0.01 before spreading onto YPD agar with a sterile cotton swab. AMB-containing discs were applied onto agar surfaces. Pictures were taken after 48 hours at 37 °C. **c, e, f**, Dose-response assays of *C. auris* strains after growing for 30 hours in RPMI at 37 °C with/without drugs as indicated (caspofungin – CAS, voriconazole – VRC, 5-fluorocytosine – 5FC, amphotericin B – AMB, acetazolamide – ACZ). **d**, MIC_100_ for AMB recorded at 48 hours in Fig. 2d (top) and 2e (bottom). **g**, Comparison of MIC_100_ in media with or without 2% glucose.

**Extended Data Fig. 4.**
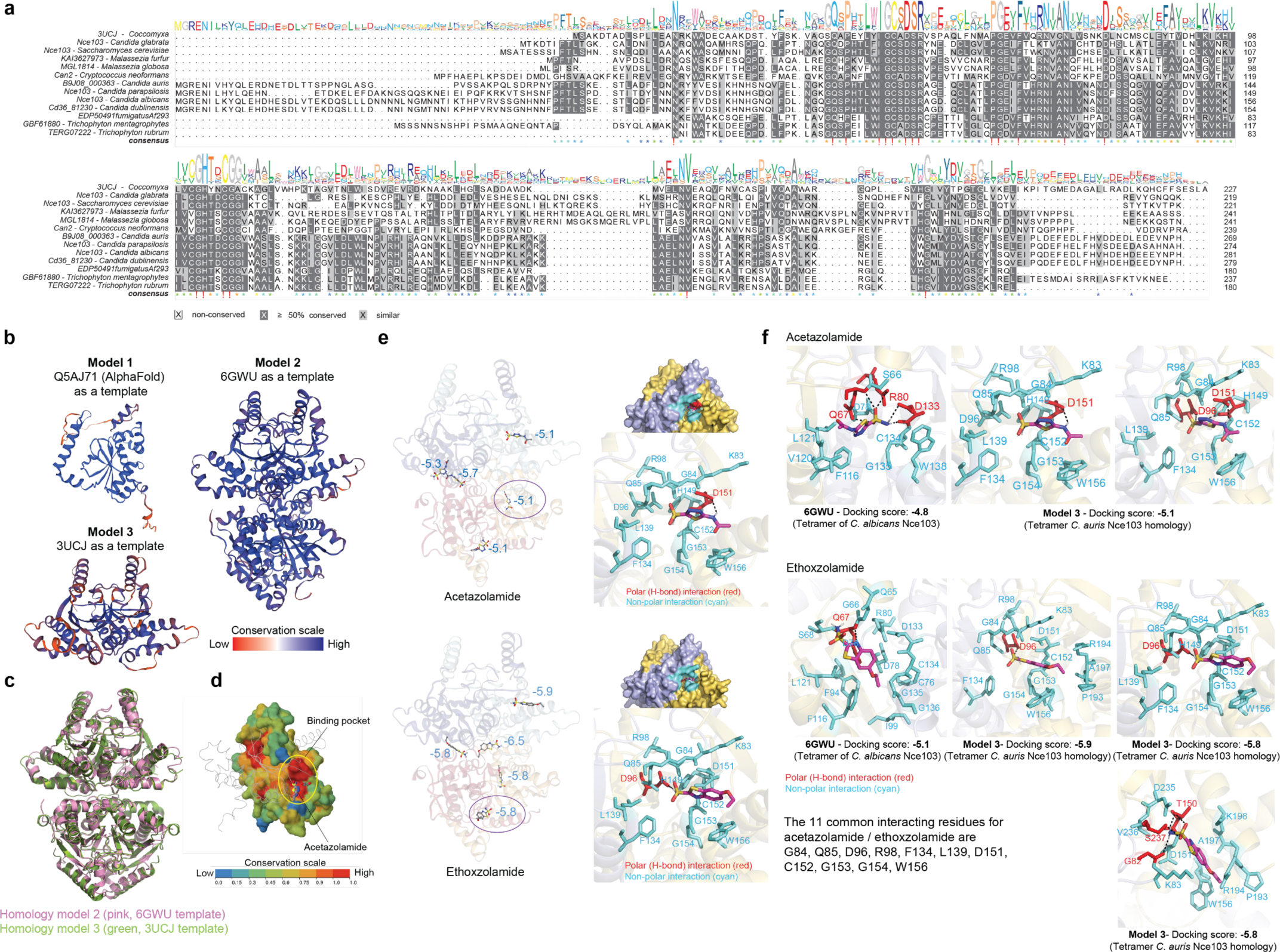
│ Nce103 is highly conserved across fungal species. **a**, Multiple sequence alignments of Nce103 homologues across fungal species. Query protein sequences were subjected to *blastp* on *C. auris* Nce103. The best hits were reciprocally searched against *C. auris*, confirming that the open reading frame B9J08_000363 encodes carbonic anhydrase (Nce103). **b**, Homology models of Nce103 generated by the Alphafold tool (Model 1), *C. albicans* Nce103 (Model 2), and Coccomyxa 3UCJ (Model 3). **c**, The overlay of models 2 and 3 yield a highly similar structural prediction. **d**, Nce103 holds a conserved binding pocket for acetazolamide in beta carbonic anhydrases. **e**, Docking results of 2 specific inhibitors of carbonic anhydrase to Model 3. **f**, Binding pocket analysis reveals 11 shared interacting residues for both ACZ and ETZ.

**Extended Data Fig. 5.**
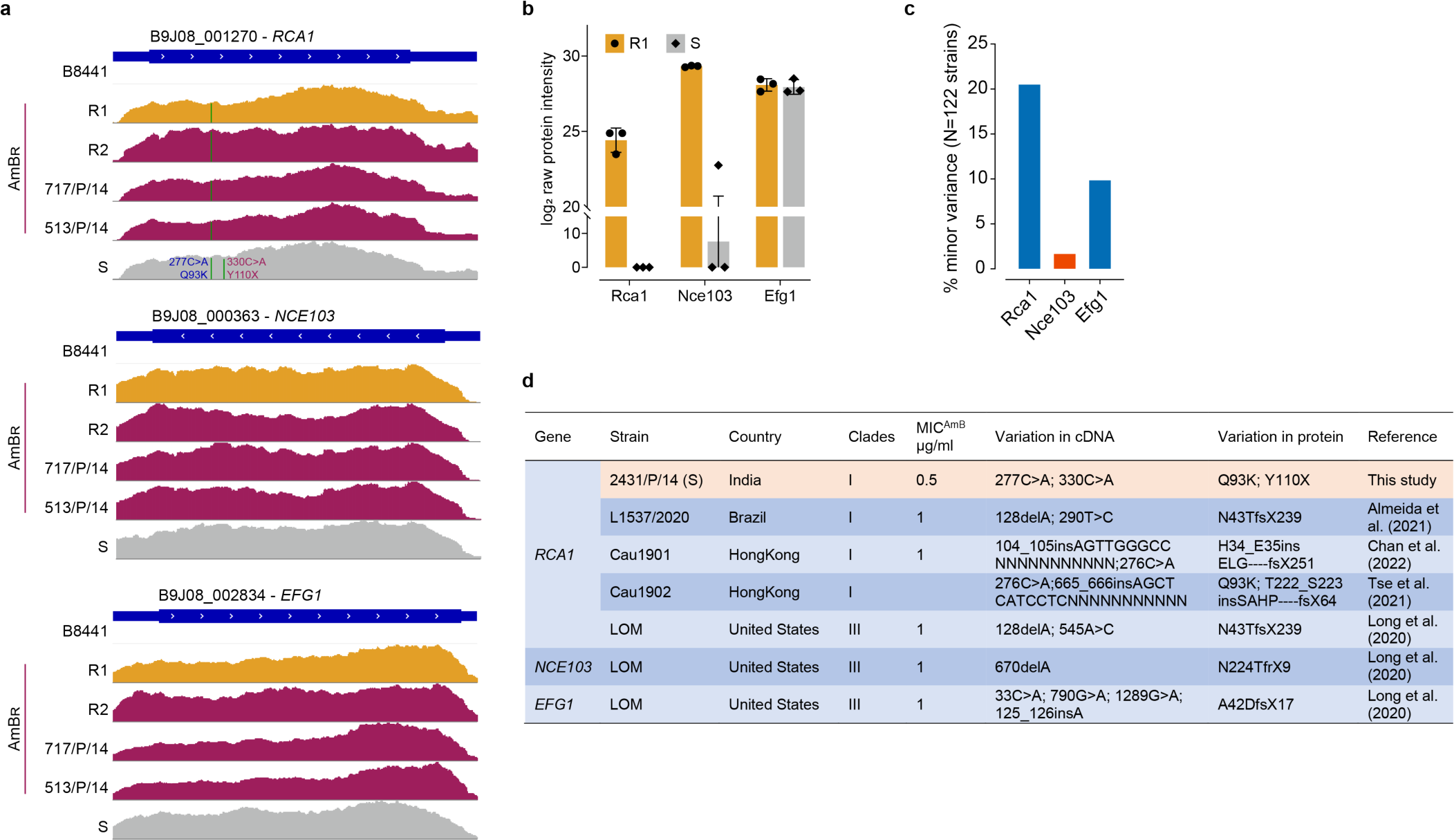
│ Mutation in key genes of the CSP correlate with amphotericin B susceptibility. **a**, IGV visualization of RNA-seq data reveals a nonsense mutation in *RCA1* sequence of the S strain. **b**, Raw protein intensity from proteomics data showed no detectable Rca1 in strain S. **c-d**, Key genes of CSP were searched with *blastn* against whole-genome shotgun contigs (wgs) and NCBI-hosted nucleotide collection (nr/nt) databases found data for *C. auris* 122 strains. Percentage of minor variances among 122 strains is shown **(c)**. Frameshift mutations led to truncated proteins being shown in **(d)**. X: stop codon.

**Extended Data Fig. 6.**
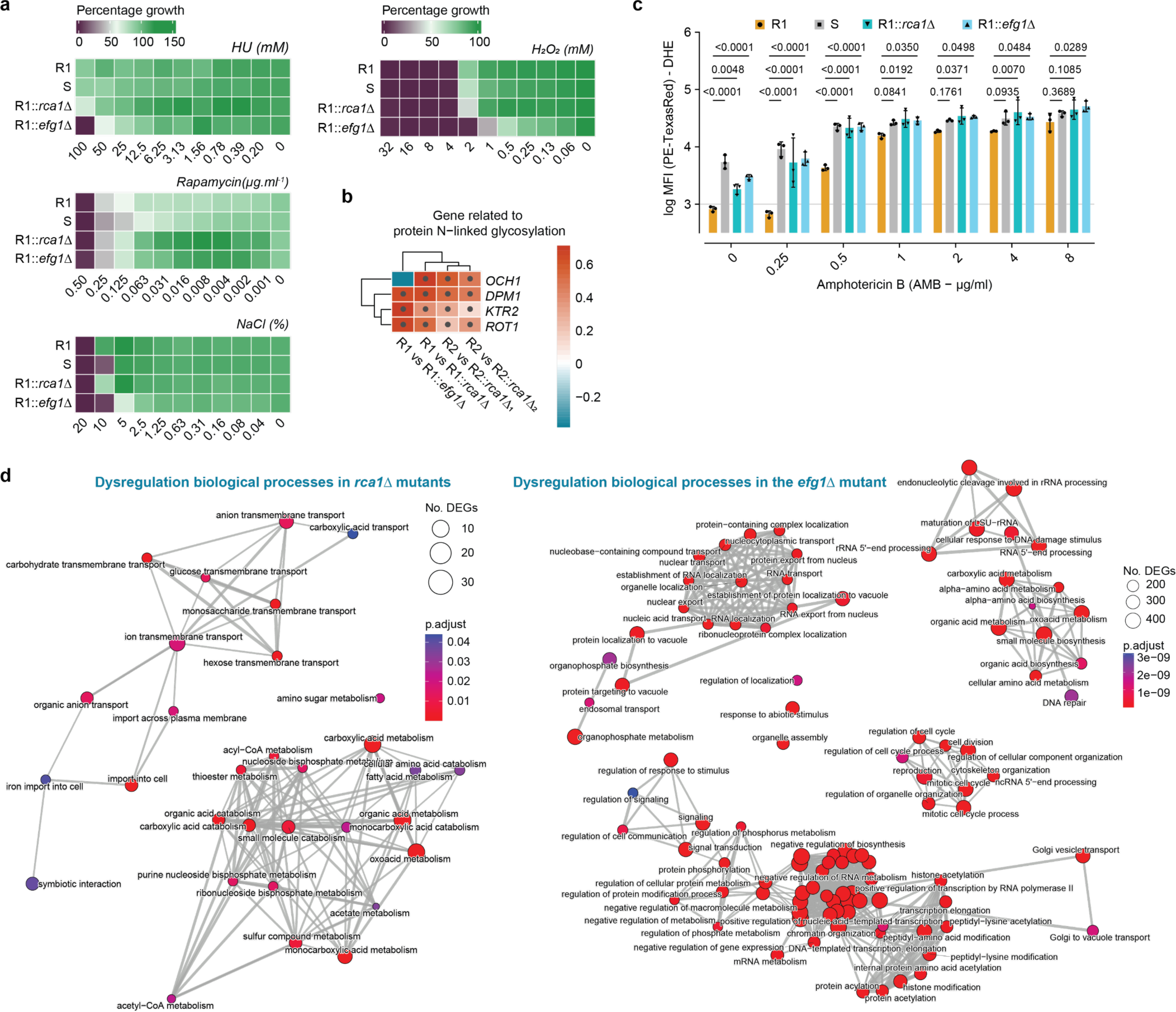
│ Stress response pathways implicated in AMB^R^ traits. **a**, Dose-response assays of *C. auris* strains growing for 30 hours in RPMI at 37 °C with or without cell wall stress agents (Calcofluor White – CFW, Congo Red – CR), mitochondrial inhibitor (Antimycin A), oxidative stressor (H_2_O_2_), osmotic stressor (NaCl), genotoxic stressor (hydroxy urea – HU) and Tor1 inhibitor (Rapamycin). **b**, DEGs related to N-linked protein glycosylation filtered with a ± 0.58 cut-off log_2_FC for *rca1*Δ. Color scale indicates log_2_FC. **c**, Intracellular ROS response to different AMB concentrations presented in Fig. 3b. Mean fluorescent intensity (MFI) of the PE-TexasRed channel were exported by FlowJo-gating for single cells. P values were calculated by two-tailed unpaired t-tests with multiple comparisons correction by BH. **d**, Similarity matrix among significant GO enrichment terms of DEGs (cut-off log_2_FC ± 0.58) in *rca1*Δ and *efg1*Δ versus WT. Interaction of enriched pathways visualized for *rca1*Δ (left), *efg1*Δ (right).

**Extended Data Fig. 7.**
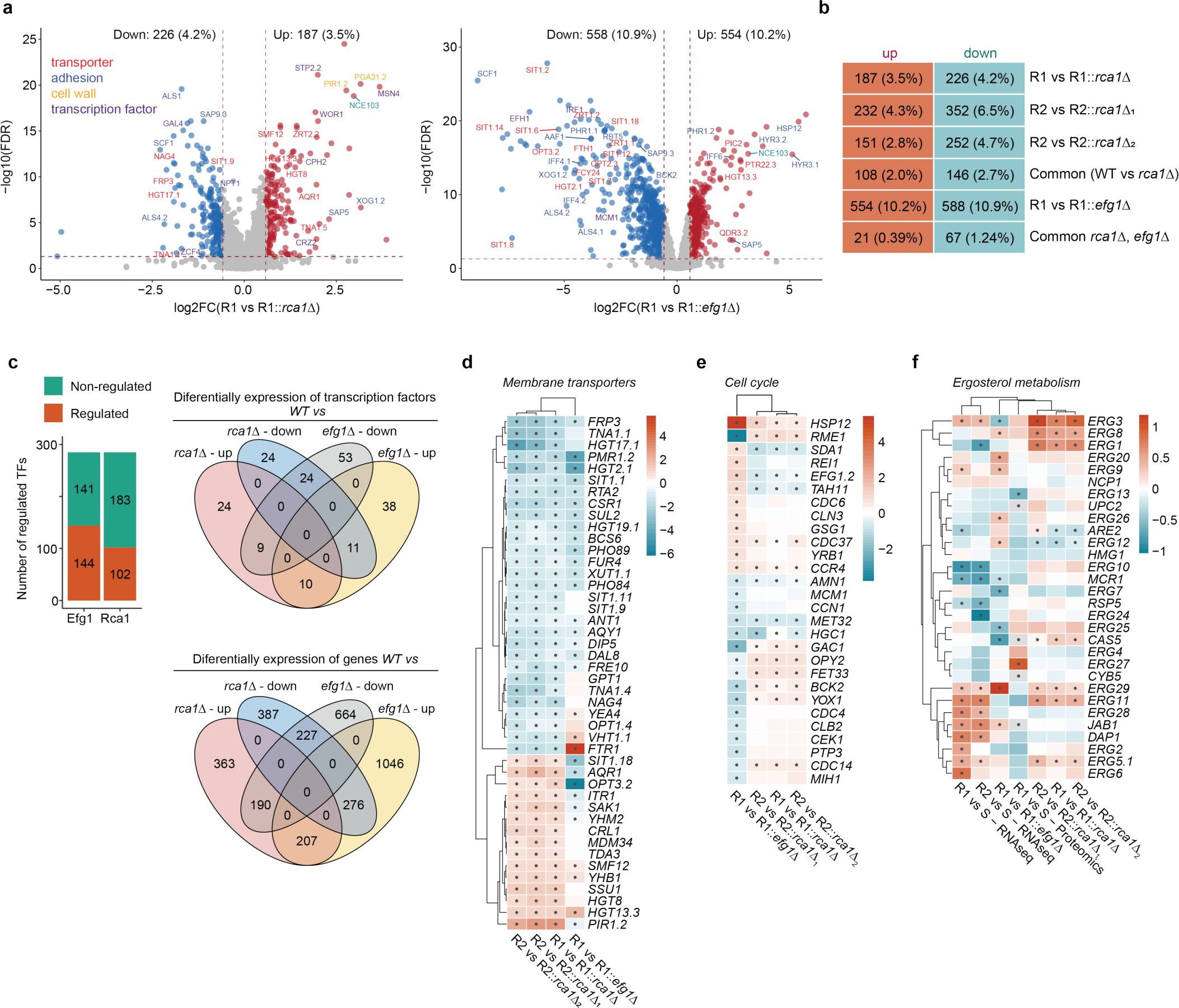
│ Global dysregulation of transcription upon ablation of Rca1 or Efg1. **a**, Representative Volcano plots of R1 versus *rca1*Δ and *efg1*Δ with cut-off log_2_FC ± 0.58, adjusted p value < 0.05. **b**, Number of up– and down-regulated genes in WT versus *rca1*Δ and *efg1*Δ. Percentage indicates percent over total number of expressed genes in RNA-seq. **c**, Number of transcription factors and DEGs regulated by Rca1 and Efg1; data shows all DEGs with adjusted p value < 0.05. **d-f**, Integrated heatmaps from different omics data sets identify genes related to ergosterol metabolism, cell cycle and membrane transporters. Color scale indicates log_2_FC. Black dots indicate adjusted p-value < 0.05.

**Extended Data Fig. 8.**
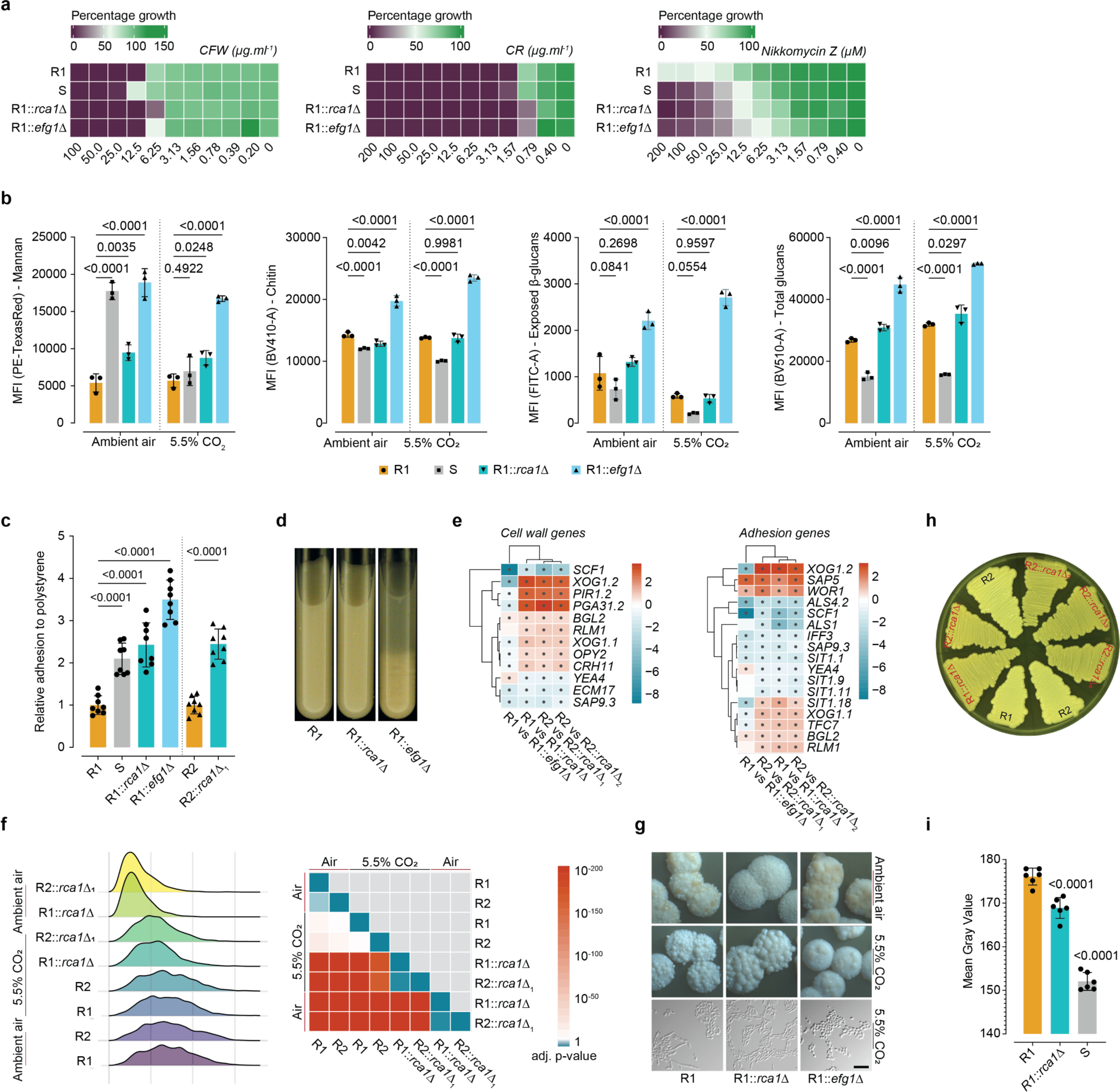
│ Rca1 and Efg1 are master regulators of morphogenesis and cell surface remodeling. **a**, Dose-response assays of *C. auris* strains after 30 hours culture in RPMI at 37 °C with or without cell wall stressors (Calcofluor White – CFW, Congo Red – CR, Nikkomycin Z-NKZ). **b**, Mannan, chitin, surface-exposed β-glucans, total glucans stained with Concanavalin A-Texas Red, CFW, Fc (human): Dectin-1 (mouse) followed by Alexa Fluor^®^ 488 anti-human IgG Fc and aniline blue, respectively, after 6 hours culture in YPD. Graph shows MFI measured by flow cytometry. **c**, Adhesion to polystyrene plastic plates by crystal violet staining. **d**, Cells were cultured for 18 hours, washed twice in PBS and adjusted to 5×10⁷ cells/ml PBS in glass tubes for 15 min flocculation assays. **e**, Heatmaps show DEGs involved in cell wall architecture and adhesion. DEGs data were extracted from RNA-seq datasets of WT strains and independent *rca1*Δ mutant clones. Data for *efg1*Δ were included after filtering with *rca1*Δ cut-off log_2_FC of ± 0.58. Color scale shows log_2_FC values; black dots indicate adjusted p-value ≤ 0.05. **f**, Cell size analysis using flow cytometry after 48 hours of culture in RPMI. FCS-A values were exported and visualized. Adjusted p-values from a pairwise Wilcox test, multiple correction by Benjamini-Hochberg (BH) were visualized in a heatmap. **g**, Colony morphology of WT and deletion strains cultured on YP-glycerol agar for 14 days at 37 °C (top); scale bar is 1mm. Cell morphology (bottom) of cells cultured in RPMI at 37 °C for 24 hours; scale bar is 10µm. **h-i**, Representative picture comparing color of WT and *rca1*Δ grown in rich YPD medium. **i**, Quantification of mean gray values of WT and *rca1*Δ colonies after 3 weeks on YPD agar. Two-way ANOVA with Dunnett’s test (**b, c, i)** and two-tailed unpaired t-test with multiple comparisons corrections by BH **(f)**.

**Extended Data Fig. 9.**
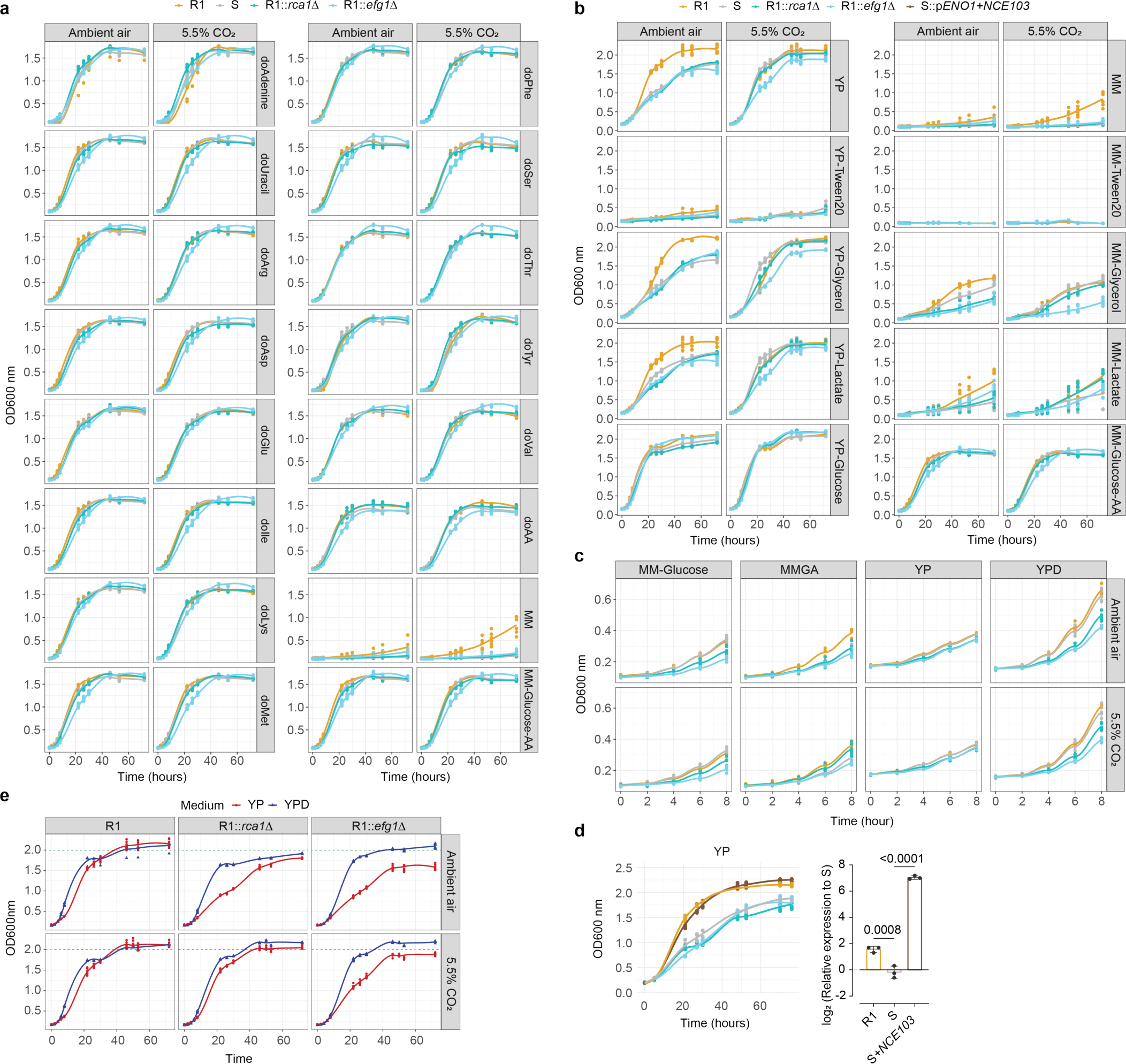
│ Energy metabolism plays important roles in fitness of *rca1*Δ *and efg1*Δ. **a-b**, Growth curves in various culture media at OD_600nm_ were recorded over 72 hours at 37 °C; **a**, drop-out (do) media with different amino acid or **b**, containing different carbon sources. **c**, Growth kinetics within the first 8 hours. **d**, Overexpression of *NCE103* in strain S improved growth similar to R1 (left). Relative expression of *NCE103* (right). One-way ANOVA with Dunnett’s multiple comparisons test was applied. MM: minimal medium, do: drop-out; AA: complete amino acid mixes. Detailed media formulations are described in Supplementary Table 2. **e**, Comparing fitness in YP and YPD highlights the critical roles of CSP and glucose in enabling *C. auris* to produce biomass.

**Extended Data Fig. 10.**
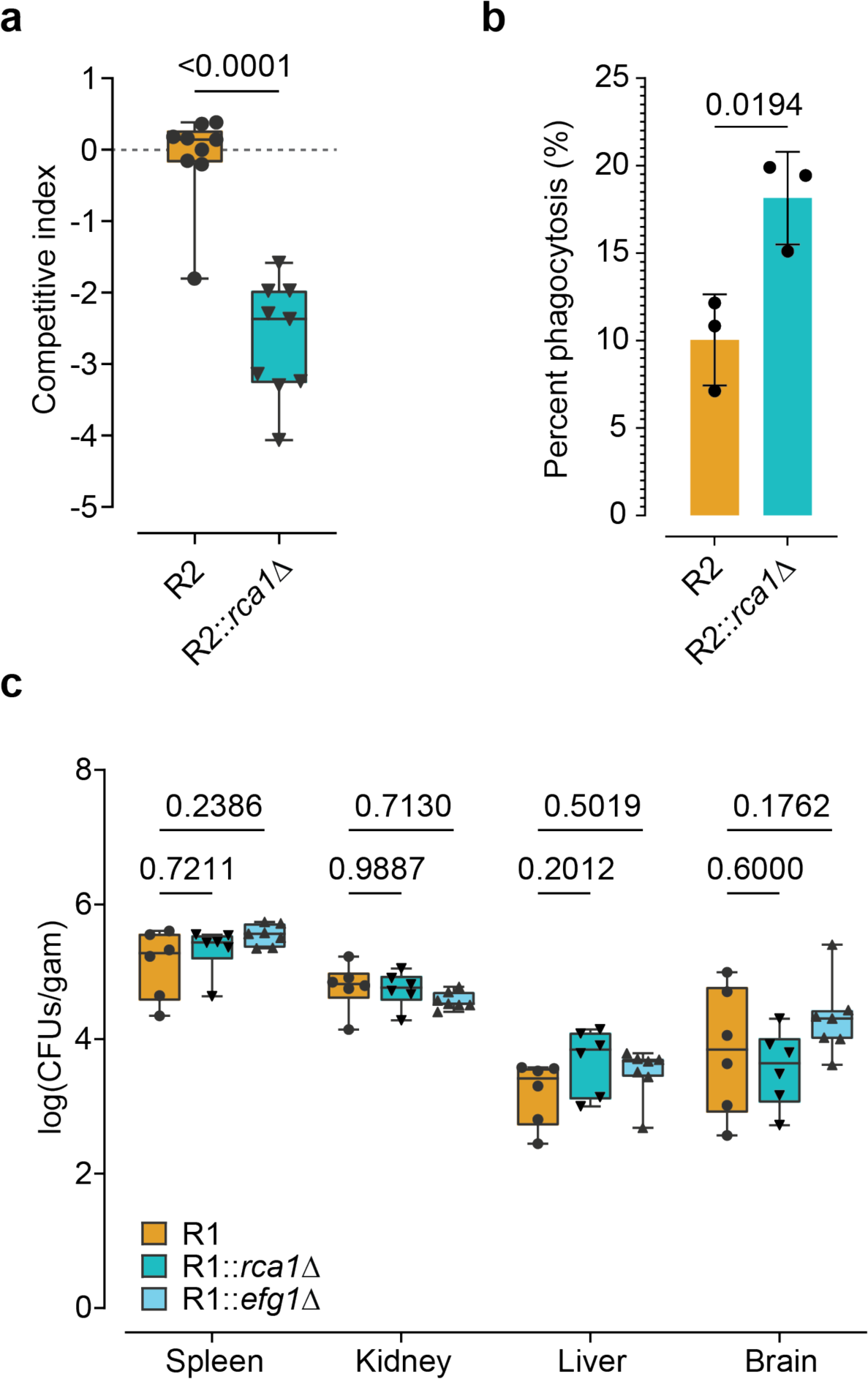
│ Fitness defects of *C. auris* lacking Rca1 *in vivo*. **a**, Competitive fitness index based on CFUs recovered from mouse back skin surfaces after a 14-day colonization period (n=9). **b**, Phagocytosis assays were conducted with primary bone marrow-derived macrophages (BMDMs). Fungal cells were labeled with FITC and subjected to BMDMs-fungal interaction for 60 minutes. Fungal phagocytosis was analyzed by flow cytometry. **c**, C57BL/6 mice were tail vein injected with 2×10⁶ fungal cells in 100 µl per 21.5 grams body weight. Fungal burden in organs was quantified at day five post infection (5-7 male mice). P values were calculated by two-tailed unpaired t-test **(a, b)** or Two-way ANOVA with Tukey’s multiple comparisons test **(c)**.

**Extended Data Fig. 11.**
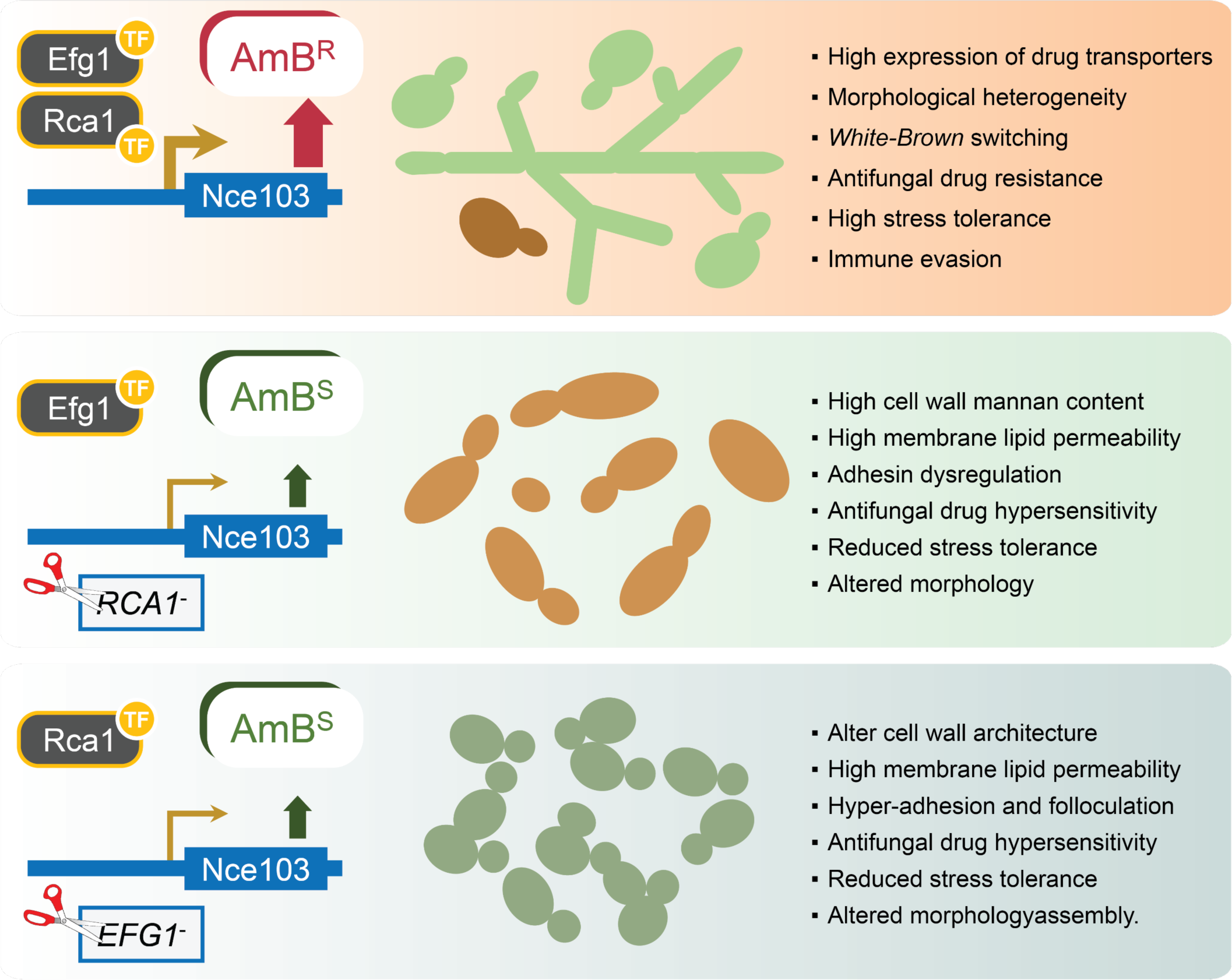
│ Cooperative functions of Rca1 and Efg1 in *C. auris* pathophysiology. WT (top), *rca1Δ* (middle), *efg1Δ* (bottom).

**Extended Data Fig. 12.**
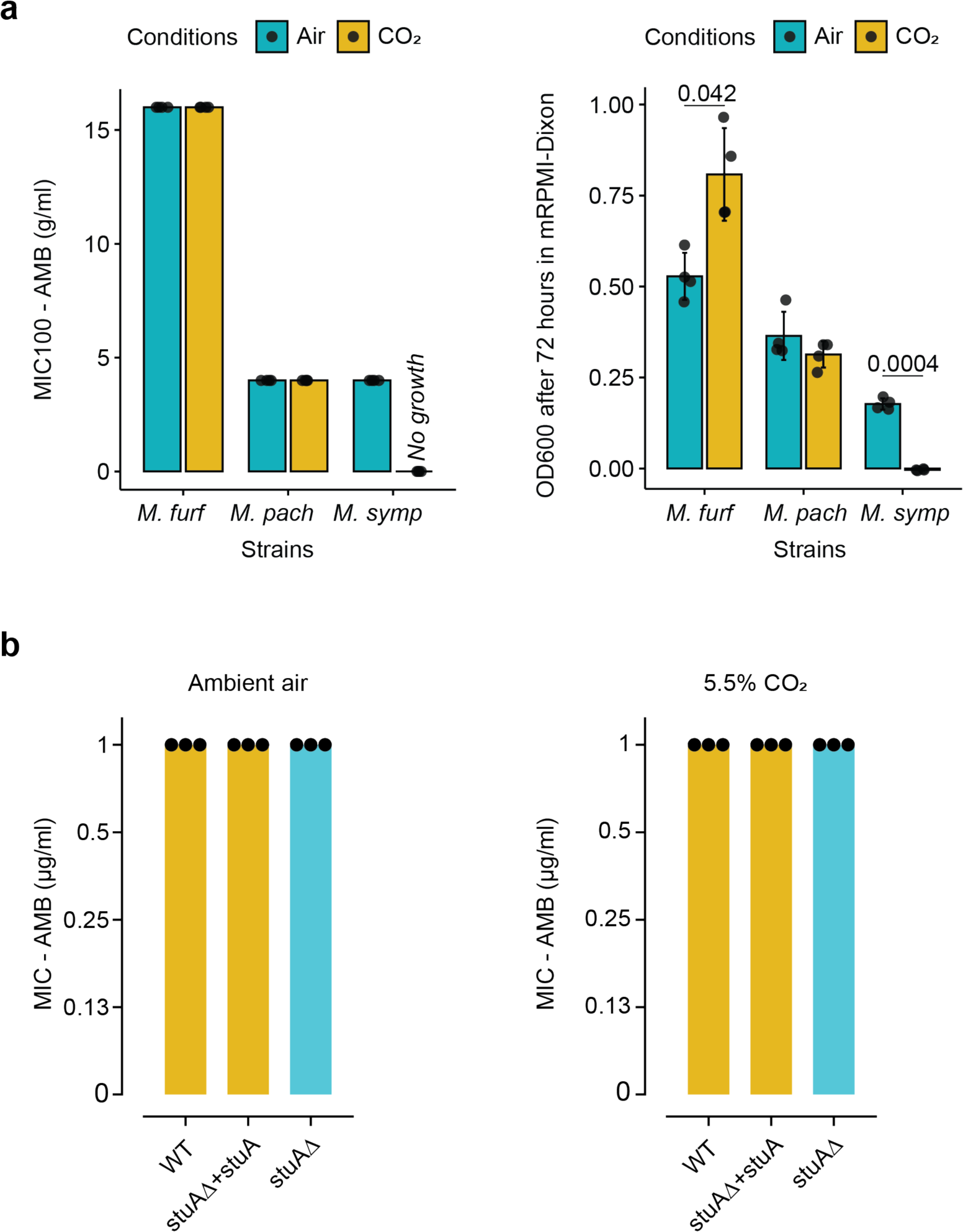
│ Malassezia spp and Dermatophytes do not employ CSP for AMB^R^. **a**, MIC_100_ values of Malasezia with AMB (left) and growth of Malassezia after 72 hours in MIC-testing medium RPMI-mDixon. *Malassezia furfur* (*M. furf*), *Malassezia pachydermatis* (*M. pach*), *Malassezia sympodialis* (*M. symp*). P values were calculated by two-tailed unpaired t-test with BH correction. **c**, MIC assay for *Trichophyton benhamiae (Arthroderma benhamiae)* and its *stuA* mutant (homologue of *EFG1*) with AMB, conducted in RPMI medium after 48 hours at 30 °C.

**Extended Data Fig. 14.**
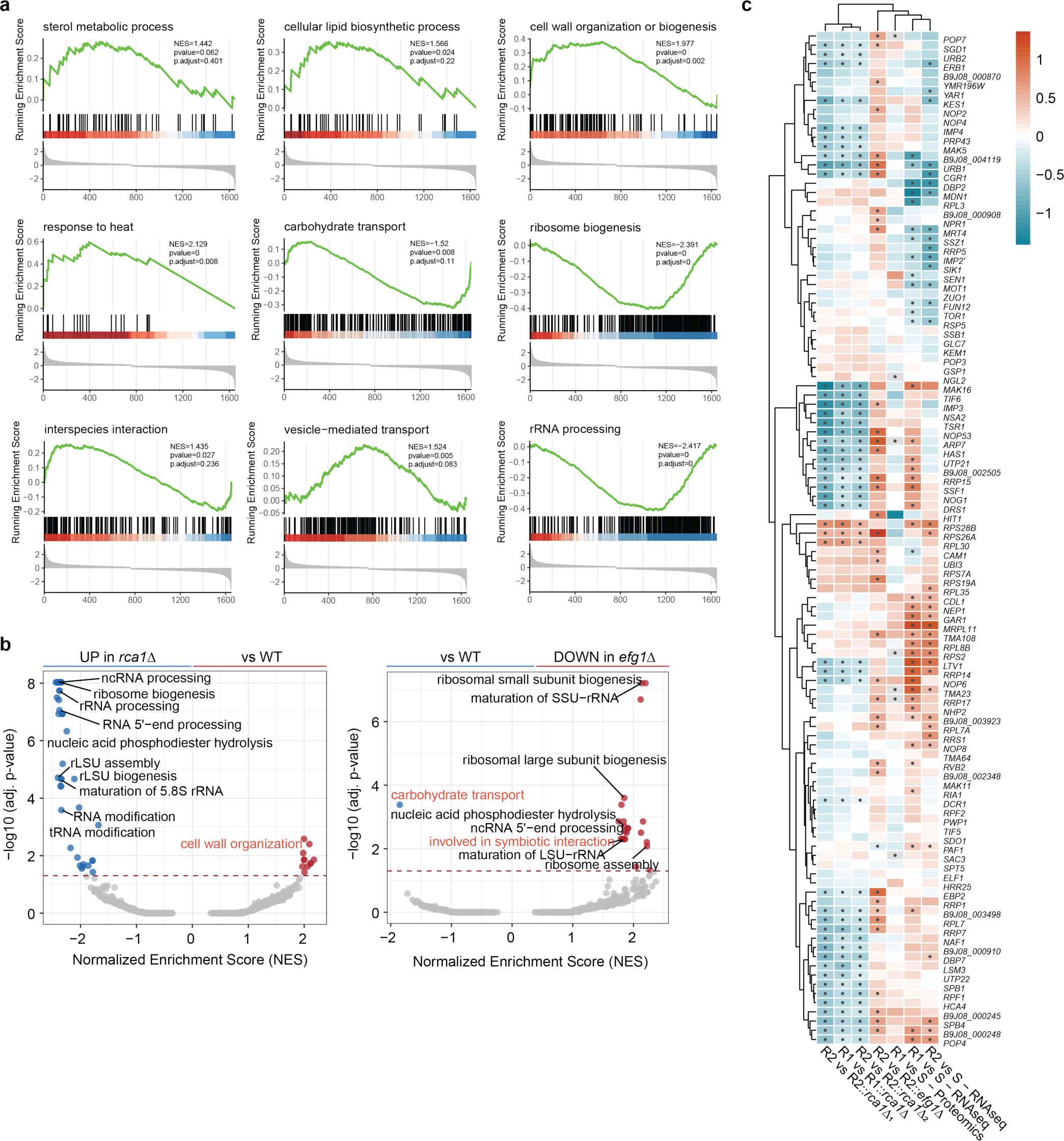
│ Gene set enrichment analysis (GSEA) of RNA-seq data for *rca1*Δ and *efg1*Δ mutants. **a**, Representative results of GSEA using gseGO (*clusterProfiler*). Positive Normalized Enrichment Scores (NES) indicate pathways enriched in WT strains. Negative NES indicate pathways enriched in *rca1*Δ. **b**, GSEA-filtered genes with adjusted p-value < 0.05. Volcano plots show GSEA data for WT, *rca1*Δ and *efg1*Δ cells; red line reflects adjusted p-value = 0.05. **c**, Heatmap showing integration of omics datasets – filtered genes related to RNA processing and ribosome biogenesis from GO biological processes; color scale indicates log_2_FC; black dots indicate adjusted p-value < 0.05.

## Supplementary information (SI)

### SI 1. Materials and methods

Detailed methods can be found in this section.

### 1.1. Dose-response assays for filamentous fungi and Malassezia spp

For filamentous fungi, fungal cells were grown on YPD agar for 5-7 days. Conidia were harvested in 2 ml of PBS and allowed to settle for 5 minutes (min). The upper suspension was transferred to the new microcentrifuge tube followed by measured with a cell counter CASY (Roche). Subsequently, cells were diluted to a concentration of 5×10^4^ conidia/ml and 100 µl of this solution was added to plates containing testing reagents prepared using the exact procedure described in Method section for dose-response assay. For Malassezia^1^, fungal cells from m-Dixon agar plates were cultivated in fresh liquid medium for 2 days. Cells were adjusted to a density of 5×10^5^ cell/ml in RPMI-m-Dixon medium before adding into a 96-well plate preparing with serial dilution of testing agents. Minimal inhibitory concentrations (MIC_100_) were visually recorded after incubation for 48 hours.

### 1.2. Spotting and disc diffusion antifungal assays

Fungal cells from overnight cultures were reinoculated from an initial 0.2 OD_600_ nm for 5 hours. Cells were counted by a cell counter CASY then adjusted to 10^7^ cells/ml in PBS. For spotting assay, a 5-fold serial dilution was performed in a 96-well plate. 0.5 µl of each dilution was spotted onto YPD agar without/with antifungal agents (different concentrations of amphotericin B and 5-fluorocytosine). Plates were incubated at 37 °C ambient air and 5.5% CO_2_. Pictures were taken after 2 days.

Disc diffusion assay was performed following CLSI M44-A^2^, a sterile cotton swab was dipped into a 10^6^ fungal cells suspension and streaking on YPD agar. Paper discs (BioRad) containing 100-unit amphotericin B were put on top. The inhibitory zone diameter was measured after 3 days at 37 °C.

### 1.3. Proteomics

Several colonies of *C. auris* growing on YPD agar were picked and re-grown overnight in RPMI 1640 (Gibco) buffered with 35 g/l MOPS (AppliChem), pH 7 (RPMI). Fungal suspensions were transferred to 50 ml fresh RPMI to reach and OD_600nm_ of 0.1 in baffled flasks. After 7 hours incubation at 30 °C with agitation of 200 x rpm, yeast cells were pelleted and washed 3 times in cold PBS (Sigma-Aldrich). Cells were resuspended in tubes containing 300 mg of glass beads (Sigma-Aldrich) and 1 ml of Candida lysis buffer (1% sodium deoxycholate – SDC, 100 mM Tris-HCl, 150 mM NaCl, 1 mM PMSF, 1 mM EDTA 1 tablet/ 50 ml complete protease inhibitor), followed by bead beating (FastPrep-MPI) 6 m/s × 45 s × 3 times. The supernatant was collected by centrifuging through the small hole created at the bottom of the tube with a G26 needle tip. Protein was precipitated by 4 volume acetones, at –20 °C, overnight.

The protein pellets were resuspended in 300 µl4% (w/v) SDS, 100 mM Tris/HCl pH 8.5, and incubated at 95 °C for 5 min. The lysate was clarified by centrifugation at 16000g for 10 min at 30 °C. The supernatant was then transferred to a new tube and protein concentrations were measured using a 600 nm protein assay kit (Pierce) with SDS compatibility reagents. 50 µg protein was reduced by adding 30 µl of 1 M dithiothreitol (DTT) and heated at 95 °C for 5 min. Samples were diluted with 9X 8 M urea in 100 mM Tris/HCl pH 8.5, transferred to the FASP filter, and centrifuged for 20 min at 12,000 g. After washing with 200 µl 8 M urea in 100 mM Tris/HCl pH 8.5 for 20 min at 120,00 g, 100 µl 50 mM iodoacetamide in 100 mM Tris/HCl pH 8.5 was added, vortex-mixed for 1 min and then incubated for 30 min in the dark at room temperature. After centrifugation for 10 min at 12,000 g, samples were washed twice with 200 µl 8 M urea in 100 mM Tris/HCl pH 8.5 and three times with 100 µl of 50 mM ammonium bicarbonate (ABC). The filter was transferred to a new collection tube and 40 µl of 50 mM ABC containing 1 µg trypsin platinum (Promega) was added and kept at 37 °C overnight. Digested peptides were collected by centrifuging for 15 min at 12,000 g. The filter was washed with 40 µl 50 mM ABC, centrifuged for 15 min at 12,000 g, and pooled. The digested peptides were acidified with 10 µl 10% TFA and the peptides were desalted using C18 Stagetips^3^ and MCX 96 well plate (Waters).

Tryptic peptides were separated on an Ultimate 3000 RSLC nano-flow chromatography system (Thermo-Fisher), using a pre-column for sample loading (Acclaim PepMap C18, 2 cm × 0.1 mm, 5 μm, Thermo-Fisher) and a C18 analytical column (Acclaim PepMap C18, 50 cm × 0.75 mm, 2 μm, Thermo-Fisher), applying a segmented linear gradient from 2 to 35% and finally 80% solvent B (80% acetonitrile, 0.1% formic acid; solvent A 0.1% formic acid) at a flow rate of 230 nl/min over 120 min. Eluted peptides were analyzed on an Exploris 480 Orbitrap mass spectrometer (Thermo-Fisher) coupled to the column with a FAIMS pro ion-source (Thermo-Fisher) using coated emitter tips (PepSep, MSWil) using the following settings. The mass spectrometer was operated in DDA mode with two FAIMS compensation voltages (CV) set to –45 or –60 and 1.5 s cycle time per CV. The survey scans were obtained in a mass range of 350-1500 *m/z*, at a resolution of 60k at 200 m/z, and a normalized AGC target at 100%. The most intense ions were selected with an isolation width of 1.2 *m/z*, fragmented in the HCD cell at 28% collision energy, and the spectra recorded for max. 50 ms at a normalized AGC target of 100% and a resolution of 15k. Peptides with a charge of +2 to +6 were included for fragmentation, the peptide match feature was set to preferred, the exclude isotope feature was enabled, and selected precursors were dynamically excluded from repeated sampling for 45 seconds.

Raw data were split into single cv using FreeStyle 1.8 SP2 and processed using the MaxQuant software package (version 1.6.17.0)^4^. Data were mapped to the Uniprot *C. auris* reference proteome (version 2021.03, www.uniprot.org), as well as a database of most common contaminants. The search was performed with full trypsin specificity and a maximum of two missed cleavages at a protein and peptide spectrum match false discovery rate of 1%. Carbamidomethylation of cysteine residues were set as fixed, oxidation of methionine and N-terminal acetylation as variable modifications. For label-free quantification the “match between runs” feature and the LFQ function were activated – all other parameters were left at default.

MaxQuant output tables were further processed in R using Cassiopeia_LFQ (https://doi.org/10.5281/zenodo.5758974). Reverse database identification, contaminating proteins, protein groups identified only by a single modified peptide, protein groups with less than two quantitative values in one experimental group, and protein groups with less than 2 razor peptides were removed for further analysis. Missing values were replaced by randomly drawing data points from a normal distribution modelled on the whole dataset (data mean shifted by –1.8 standard deviations, width of distribution of 0.3 standard deviations). Differences between groups were statistically evaluated using the LIMMA package^5^ at 5% FDR (Benjamini-Hochberg).

### 1.4. RNA isolation

Fungal cultures were centrifuged, and cell pellets were rapidly frozen in liquid nitrogen. Dry cell pellets were then stored at –80 °C for later use. For RNA isolation, the fungal cells were resuspended in 1 ml of TRI Reagent (LabConsulting) and disrupted by two times FastPrep bead-beating (MP Biomedicals) with 300 mg glass beads at 6 m/s for 45 seconds. Samples were centrifuged at 14,000 g for 10 min at 4 °C. The aqueous phase was further purified by adding 200 µl of chloroform, followed by precipitation with isopropanol at –20 °C for 30 min. RNA pellets were collected by centrifugation at 14,000 g for 20 min at 4 °C, pellets were washed with cold 70% ethanol before air-drying the RNA. Finally, RNA pellets were resuspended in 25 µl RNAse-free water (Invitrogen).

Each sample equivalent to about 5 µg of total RNA, underwent treatment with RNAse-free 10U of DNase I (Thermo Scientific) for 15 min in the presence of 50U of RiboLock RNase Inhibitor (Thermo Scientific). Subsequently, the RNA was purified by PCI extraction. The aqueous phase containing RNA was collected by ethanol precipitation with 30 mM sodium acetate at pH 5.3. RNA quality and purification was checked with Nanodrop and conventional PCR targeting *ACT1*.

### 1.5. Genomic DNA isolation

Fungal cultures were centrifuged, and cells washed 2 times with sterile distilled water, followed by phenol – chloroform – isoamyl alcohol (PCI) extraction method. Cells were resuspended in Yeast Lysis Buffer (2% Triton X-100, 1% SDS, 100 mM NaCl, 1 mM EDTA, 10 mM Tris-Cl pH 8). The same amount of PCI (Sigma-Aldrich) was added before cell disruption by bead-beating (FastPrep-MPI) 6 m/s x 45 s x 2 times. Released DNA was washed with chloroform then precipitated from the aqueous phase with ethanol and acetate. DNA pellets were resuspended in water then treated with DNAse-free RNase A (Thermo Scientific – EN0531). DNA was reprecipitated with 40 mM ammonium acetate in ethanol at –20 °C and dissolved in nuclease free water (Invitrogen – 10977035).

### 1.6. Integration of transcriptomics and proteomics datasets

A *C. auris* annotation database was generated by integrating annotation data from the Candida Genome Database, FungiDB and UniprotKB. *C. auris* protein metadata (taxonomy_id:498019) were retrieved from Uniprot as a tsv file. Protein ID, gene ID, protein names were extracted. Homology data files of *C. auris* with other yeast species were retrieved from CGD. Gene Annotation File (.gaf) released version 54 of *C. albicans* SC5314 were retrieved from FungiDB^6^. All data sets were merged into a unique data frame based on gene ID and protein ID. This annotation database (AnDB) was used for all analysis with some manual curation.

For published transcriptomic data, we retrieved our published datasets from the same *C. auris* strains from Gene Expression Omnibus (GEO), including GSE198410 and GSE190920. Raw read counts from those datasets were used for DEG analysis with EdgeR. Differential abundance analysis from proteomics experiments were merged with RNA-seq DEGs results in the same data frame based on protein and gene ID before mapping with AnDB. For integrative analysis, data were filtered using *str_detect* for certain gene ontology, protein name and descriptions followed by visualization using the pheatmap^7^, VennDiagram^8^, and ggplot2^9^ packages. For gene set enrichment analysis (GSEA) and gene ontology (GO) analysis, specific *C. albicans* geneID (assembly 22 ID) were used as input in gseGO or enrichGO functions from clusterProfiler^10^ using an GO database constructed from a *gaf* file v54 of *C. albicans* SC5314. Results from those analysis were visualized by enrichplot^10^.

### 1.7. Quantification of cell wall carbohydrate components

Fungal cells from overnight cultures were inoculated into fresh YPD medium to an initial OD_600nm_ of 0.2, and incubated for 6 hours at 37 °C under 5.5% CO_2_ or ambient air. Cells were quantified using the CASY Cell Counter. Next, 2×10^6^ cells were washed twice with FACS buffer and pelleted for cell wall carbohydrate staining as described before^11^. Exposed mannan was stained by incubating cells for 45 min at 30 °C in 1 ml FACS buffer containing 25 μg/ml Concanavalin-TexasRed (Invitrogen). To stain exposed β-glucans, cells were treated on ice for 60 min with 5 ng/µl Fc:Dectin-1 (AdipoGen) in 200 µl FACS buffer. Cells were then washed for 60 min with FACS buffer, and then treated for 45 min with a 2.5 μg/ml solution of Alexa Fluor 488 anti-human IgG Fc antibody (BioLegend). For chitin staining, we treated cells with 25 µg/ml CFW in FACS buffer for 5 min. To determine the total β-glucan levels, 2×10^6^ cells were resuspended in a solution comprising 1 M glycine (pH 9.5) and 0.005% aniline blue, followed by a 5-min staining period. As controls, unstained cells were incubated for 5 min in 1 M glycine (pH 9.5). Finally, samples were analyzed using a BD-Fortessa instrument. All staining was conducted in the dark. Raw data were analyzed with the FlowJo v10.8 software using the same gating strategy described before^11^.

### 1.8. Plastic adhesion and flocculation assays

To inspect adhesion properties*, C. auris* strains from overnight cultures were grown for 4 hours in YPD at 30 °C starting from an OD_600nm_ of 0.1. Cells were washed twice in PBS and counted in a CASY Cell Counter (Roche). Cells suspensions were adjusted to 2×10^6^ cells/ml in RPMI medium, and transferred into a tissue culture 96-well plate (Starlab). Plates were incubated at 37 °C for 3 hours in a static incubator, followed by biomass quantification using crystal violet staining. Briefly, the RPMI medium was removed by inverting the plate and tapping it on towel paper. The wells then washed 2 times with PBS and fixed with 100 µl methanol (Merck) for 15 min. Plates were left to dry in a chemical safety cabinet for about 5 min, followed by a 5-min staining with 100 µl of a 0.1% aqueous crystal violet solution (Sigma-Aldrich). After three washings with distilled H_2_O, 100 µl of 33% acetic acid was added, and plates were shaken for 1 min at 800 rpm on a MixMate (Eppendorf) to extract crystal violet from cells. Absorbance was measured at OD_570nm_ in a Victor Nivo plate reader.

To visualize flocculation phenotypes, fungal cells were cultured overnight in YPD medium at 37 °C with agitation at 200 rpm, followed by washing twice with PBS. Cells were counted in a CASY cell counter, and then adjusted to 5×10^7^ cells/ml in PBS. Aliquots of 3 ml cell suspensions were transferred into test tubes and vigorously vortex-mixed for about 10 seconds before placing in racks. Flocculation was observed after settling for 15 min.

### 1.9. Fungal colony, cell morphology and color assay

Fungal cells from overnight cultures were washed 3 times with PBS. The cell number was quantified in a CASY Cell Counter (Roche), followed by plating about 200 CFUs on agar plates containing different carbon sources. Colonies morphologies were inspected and recorded with a Zeiss Axio/Stereomicroscope after 14– and 30-day incubation at 37 °C.

For assessing cell morphology, around 200 cells were inoculated into RPMI medium in 6-well plates. After 24 hours, cells were fixed with 4% paraformaldehyde (PFA). Differential interference contrast (DIC) pictures were taken with a ZEISS Axio Imager 2 microscope. For morphology quantification, cell populations were washed 3 times in cold FACS buffer (1% fetal calf serum (FCS – Gibco), 0.5 mM EDTA, 0.1% Tween-20 in PBS – both from Sigma-Aldrich), followed by analyzing with a BD-Fortessa flow cytometry. FSC-A data from single cell populations were exported and visualized with ggplot2^9^. Fungal strains were streaked on the same YPD agar plate and incubated at 30 °C for 2 weeks. Pictures of plates were taken with a V&P Scientific Imager (V&P Scientific). Mean gray values were measured from pictures using the Image J v1.53 tool.

### 1.10. Sterol lipid quantification

Fungal cells were cultured for 8 hours at 30 °C with shaking at 200 rpm, followed by inoculation into 50 ml YPD at an initial OD_600nm_ of 0.05. After 12 hours, cells were harvested by centrifugation at 3000 rpm in 3 min, resuspended in 50 ml fresh 37 °C pre-warmed YPD with/without 0.75 µg/ml amphotericin B. Flasks were incubated for 4 hours at 37 °C, with shaking at 200 rpm. Cells were pelleted at 3000 rpm for 3 min, and washed 2 times with ice-cold PBS, before flash-freezing in liquid nitrogen and lyophilizing in a freeze dryer (LSL Secfroid – Lyolab BII, Switzerland). About 5 mg lyophilized powder from each fungal sample was transferred to a 2 ml safe lock tube containing a single metal bead (3 mm diameter, Qiagen 69997) and homogenized in a bead mill (Mixer Mill MM 400, Retsch) at maximum speed for 2 min. After resuspending in 1 ml of 2 M aqueous NaOH, samples were transferred to a 4 ml glass vial and incubated at 70 °C for 1 hour, with 10 seconds of vortex-mixing every 15 min. Vials were allowed to cool at room temperature to reach about 55 °C before transferring into 2 ml microcentrifuge tubes containing 650 µl methyl *tert*-butyl ether (M*t*BE, Roth – ROT.T175). We added 100 µl internal standard (IS) cholestane (Merck – C8003) to each tube. Tubes were first shaken vigorously by hand for 1 min before centrifugation at 10000g for 5 min at room temperature. The upper layer (roughly 550 µl) was transferred into a new microcentrifuge tube containing 35 mg Na_2_SO_4_ (Merck – 238597) and 5 mg PSA (Agilent – 5982-8382). The rest of cell lysate were extracted for the second time with another 750 µl of M*t*BE as in the first extraction step. Supernatants were combined, before obtaining a clean extract by centrifugation at 10,000 g for 5 min and transfer into a glass GC vial. Solvents were evaporated overnight in a chemical hood, and redissolved in 700 µl M*t*BE before adding 50 µl of a silylation reagent mixture 10:1 of *N*-methyl-*N*-trimethylsilyl-trifluoroacetamide (MSTFA; Macherey-Nagel – 701270.201) and *N*-trimethylsilyl-imidazole (TSIM; Macherey-Nagel, – 701310.201). The samples were analyzed in a Agilent 7820A gas chromatograph (GC) coupled to an Agilent quadrupole 5977B mass spectrometer (MS). Specific sterols and precursors were identified as their corresponding trimethylsilyl (TMS) ethers by mass spectra and relative retention times (RRT) ^12^. The base peak of each sterol TMS ether was taken as a quantifier ion for determining the peak area for cholestane (IS) *m/z* 217 RRT 1.00, ergsta-5,8,22-trien-3β-ol (lichesterol) *m/z* 363 RRT 1.29, cholesta-8,24-dien-3β-ol (zymosterol) *m/z* 351 RRT 1.30, ergosta-5,7,22-trien-3β-ol (ergosterol) *m/z* 363 RRT 1.32, ergosta-7,22-dien-3β-ol *m/z* 343 RRT 1.34, ergosta-8,24(28)-dien-3β-ol (fecosterol) *m/z* 365 RRT 1.36, ergosta-5,7-dien-3β-ol *m/z* 365 RRT 1.40, ergosta-7,24(28)-dien-3β-ol *m/z* 343 RRT 1.40, ergost-7-en-3β-ol *m/z* 472 RRT 1.41, 4,4,14-trimethylcholesta-8,24(28)-dien-3β-ol (lanosterol) *m/z* 393 RRT 1.43, and 4-methylergosta-8,24(29)-dien-3β-ol (4-methylfecosterol) *m/z* 379 RRT 1.46. The sum of all detected peak areas of each sample was set as 100% and the percentage of each sterol was calculated^13,14^.

**SI 2.**
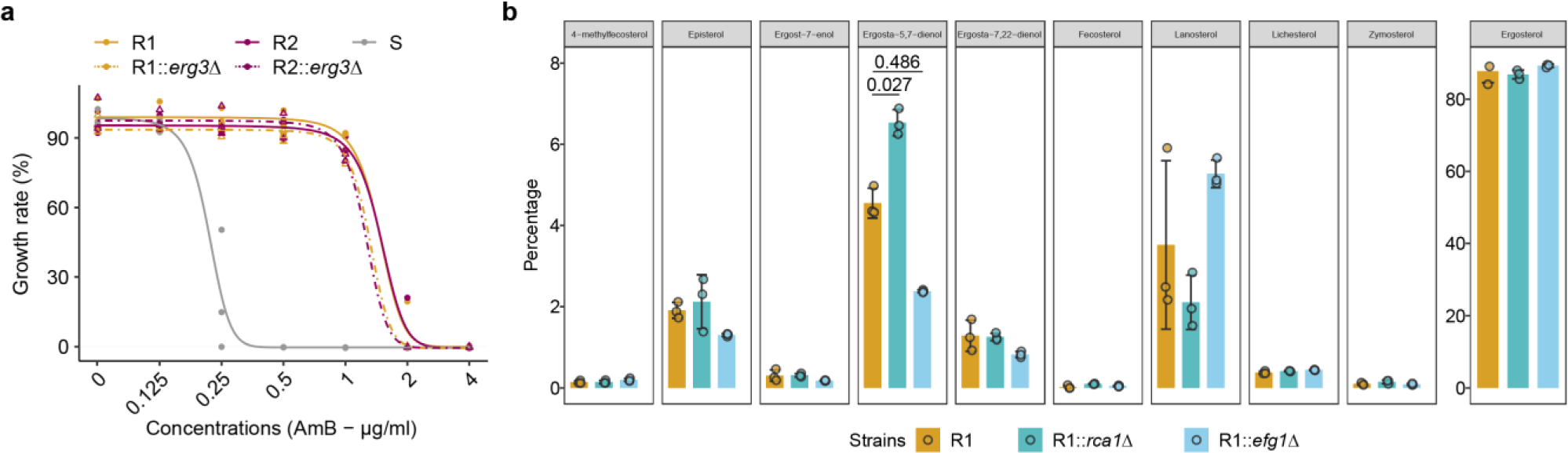
CSP modulated AmB^R^ in *Candida auris* independent with the ergosterol alteration. **a**, Deletion of *ERG3* in R strains did not affect amphotericin B resistance. **b**, Null mutants in *RCA1* or *EFG1* did not significantly affect ergosterol content in *Candida auris*.

